# General solution to inflated type I and II errors in multi-subject single-cell differential gene expression analysis

**DOI:** 10.1101/2023.01.09.523212

**Authors:** Hanbin Lee, Buhm Han

**Affiliations:** Department of Medicine, Seoul National University College of Medicine, Republic of Korea; Department of Biomedical Sciences, BK21 Plus Biomedical Science Project, Seoul National University College of Medicine, Seoul, Republic of Korea; Interdisciplinary Program in Bioengineering, Seoul National University, Seoul, Republic of Korea; Genealogy Inc

## Abstract

Large-scale multi-subject single-cell data have become very common. For differential gene expression (DGE) analysis of these datasets, a common practice is to choose any of the two: pseudobulk methods or cell-wise methods. However, multi-subject single-cell studies have highly heterogeneous study designs. Some studies have case/control samples to be compared and tested, and some make within-sample perturbations. In this work, we report that to prevent severely inflated errors, we must treat the two categories of study designs differently in the DGE analysis. In studies with case/control labels, pseudobulk methods work the best, and cell-wise methods produce severe inflation of type II errors. Cell-wise methods work best in studies with within-sample perturbations, and pseudobulk methods produce severe inflation of type I errors. We provide mathematical proofs to support this argument. Surprisingly, many existing studies, even published ones, often choose an inappropriate DGE method. The most common temptation is to use cell-wise methods for case/control studies, which will only make p-values falsely look significant. Our analyses and proofs warn that choosing an appropriate DGE method is not an option but a requirement.

## Introduction

Single-cell RNA sequencing (scRNA-seq) allowed the detection of cell-type specific differential expression patterns previously obscured in bulk RNA sequencing^1,2^. Now, multiple modalities are measured simultaneously in a single experiment^3^. Furthermore, massively parallel perturbation based on CRISPR technology is applied at a single-cell resolution^4^. Also, data are collected from both healthy and diseased individuals^5,6^.

Nevertheless, these complexities raise a challenge in differential gene expression (DGE) analysis, which compares expression levels between prespecified groups of cells in different states^1,2^. Previous studies have warned of the danger of false discoveries due to the complicated data-generating process of scRNA-seq data^7,8^. One notable phenomenon is pseudoreplication bias, which emerges in multi-subject studies. Characteristics of cell donors (or experimental conditions in which the cells were obtained) can affect the expression level of cells, causing cells within a subject to be correlated. Pseudobulk and mixed models have been proposed as solutions based on simulations and empirical findings. However, it is still unclear under which conditions pseudoreplication is problematic and needs to be corrected^7,8^.

In this work, we show that how the cell state (or group label) is determined is a crucial factor for determining the appropriate DGE analysis strategy. There are two large categories of study designs in single-cell DGE analysis: the case-control design and the within-sample perturbation design. In case-control designs where cells from healthy controls and diseased individuals are compared, the cell states of the cells are completely determined by the case-control status of the donors. On the contrary, the states of the cells vary within a donor in cases like perturb-seq experiments. For example, a commonly used dataset of peripheral blood mononuclear cells (PBMC) for the DGE benchmark has varying cell states within a donor as cells from a subject were randomly treated with interferon^9^.

Pseudobulk remains reliable in the former design, where the state of their donor completely determines the cell state. However, pseudobulk becomes overtly conservative in the latter design, where cell states are determined at a cell level. By contrast, cell-wise methods remain reliable in the latter design but become unreliable in the former design. We demonstrate these points through realistic simulation based on multi-subject datasets along with mathematical proofs. We found that mixed model remains valid in both designs as long as the sample size is sufficient but is the slowest.

Surprisingly, many studies in literature chose the analysis strategies crossly, applying pseudobulk to the latter design or cell-wise methods to the former design, which can lead to uncalibrated type 1 and 2 errors. We expect that for future researchers, the strongest temptation will be to apply cell-wise methods to case/control designs. This practice will only make the p-values falsely look significant, and must be avoided at all cost.

Finally, we analyzed two publicly available experimental perturbation datasets^9,10^. When appropriately using mixed models and cell-level tests, we obtained more signals compared to using pseudobulk, which would be an inappropriate choice for this study design. As a concluding remark, we provide a simple guide for users willing to perform differential expression analysis in single-cell data.

## Results

### Overview

We argue that what is currently called single-cell DGE analysis is a heterogeneous mixture of completely different situations that should be dealt differently. Broadly, single-cell DGE analysis can be divided into two large categories. In the first scenario, the comparison groups are assigned at the subject level. Case-control DGE analysis is an example in this category where cells from diseased and healthy individuals are compared. The second scenario assigns the group label at the cell level. In experimental perturbation studies, CRISPR or chemical perturbations are applied to individual cells, making variation of cell state within a subject^4,10,11^.

The appropriate method for detecting differentially expressed genes (DEGs) depends ultimately on which category the study belongs to because the two scenarios have completely different data-generating processes. In scenario 1, a subject is sampled with a case/control label, and the cells are subsequently obtained from the sample. Hence, the comparison group (which is the case/control label) of cells is determined prior to the sampling of cells. By contrast, in scenario 2, the comparison group is assigned at the cell level after the subjects are obtained (e.g. CRISPR affected cells/unaffected cells).

These different data-generating processes naturally lead to different scales of standard errors of the effect size estimate (log fold change). Standard error is the square root of the estimator’s variance, provided the data-generating process is repeated infinitely many times. Let *n* be the number of subjects and *N* be the total number of cells (*n* « *N*). In **Theorem 1 of Supplementary Note**, we show that under scenario 1, standard error is 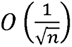, while under scenario 2, standard error is 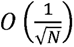. The smaller standard error in scenario 2 is intuitive, as the assignment of the comparison group is performed at a higher resolution (cell-wise).

In practice, we don’t have access to the true variance of the estimator because the experiment is done only once. Instead, a DGE method estimates the variance using the data. The problem is that this estimate can be incorrect. With few exceptions like the mixed model, most statistical tests assume independence between the observations. As a result, pseudobulk always estimates the standard error as 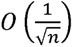 because the input data has *n* observations. Therefore, applying pseudobulk to scenario 2 leads to deflated type 1 errors and inflated type 2 errors. In contrast, cell-wise methods always approximate the standard error as 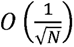. Therefore, applying cell-wise methods to scenario 1 leads to inflated type 1 errors and deflated type 2 errors. This inflation of type 1 error was already reported elsewhere^2,7,8^. We confirmed these expected phenomena by both simulations and real data analysis.

There is a lot of confusion in single-cell DGE literature. Some studies applied pseudobulk to scenario 2^10,12,13^, and some studies applied cell-wise methods to scenario 1^2,7,8,14^. Some studies simply describe the method name without any context^12^. For example, a widely used DGE method is glmGamPoi^15^, and this can be applied both via pseudobulk and cell-wisely. Thus, stating that the study used glmGamPoi does not provide enough information on how the DGE test was conveyed.

Pseudobulk is frequently used in multi-subject scRNA-seq studies in the fear of pseudoreplication bias. Traditionally, pseudoreplication bias refers to the high false discovery rate of cell-wise methods applied to multi-subject studies^2,7,8^. We show that not all multi-subject studies are subject to the bias because the bias only happens in the first scenario (**Corollary 1 and 2 of Supplementary Note**). Subjects, not cells, are the true replicates in the first case, so pseudobulk is an effective DGE analysis method. Nevertheless, in the second case, cells are the true replicates, and pseudobulk is a suboptimal choice.

Our article offers theoretically grounded but practical guidance on which method is appropriate for which situation (**Figure 1**). Pseudobulk is appropriate for scenario 1 only. Cell-wise methods are suitable for scenario 2 only. Mixed model (NB GLMM) has an interesting position in that its type 1 and 2 errors are calibrated in both scenarios as long as the sample size is sufficient. However, NB GLMM is the slowest method of all methods compared and often fails to converge for low-expression genes. In the following sections, we present simulation results that support our guidelines.

**Figure 1.**
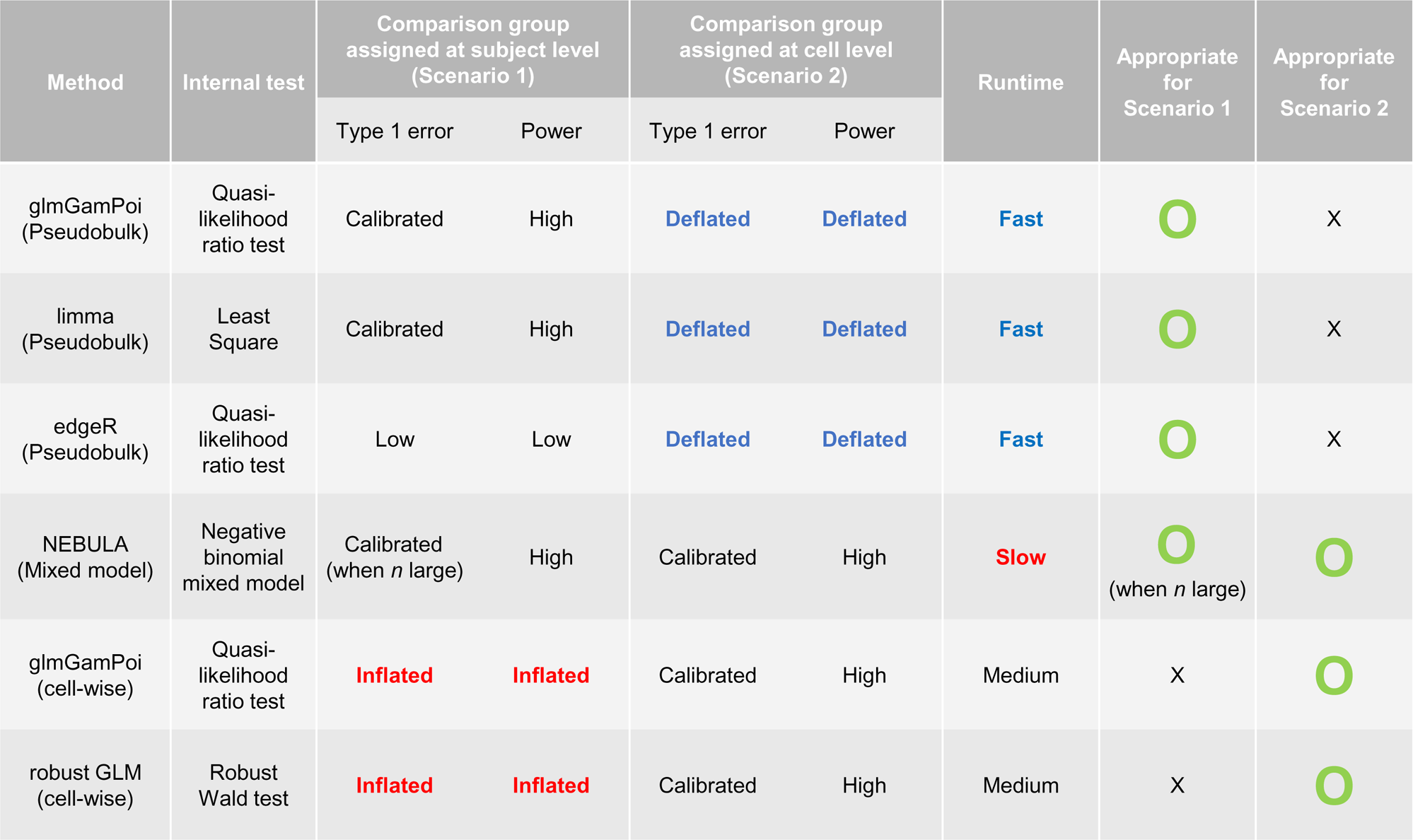
Recommendation of methods according to the study design.

### Performance of single-cell DGE methods under simulations of two scenarios

We conducted null and power simulations to evaluate the performance of the methods. Simulations were conducted for both scenarios of comparison group assignment (subject-wise and cell-wise). We compared three pseudobulk methods (glmGamPoi with aggregated counts, edgeR, and limma), two cell-wise methods (robust GLM, glmGamPoi with individual cells), and one negative binomial mixed model (NB GLMM) (see **Methods**).

An ideal *P*-value follows a uniform distribution under the null hypothesis. This means that when identical experiments are repeated, the overall distribution of the resulting *P*-values should follow a uniform distribution. Quantile-quantile (QQ) plot is a convenient way to see if a collection of *P*-values follows a uniform distribution. In the plot, randomly drawn values from the uniform distribution and the test’s *P-*values are placed in the *x-y* plane after ordering. The points are then found in the *y=x* line if the *P-*values follow the uniform distribution. Points above the line indicate that *P*-values are larger than expected, which means that the test is conservative. When the points are below the line, it means that the test is anti-conservative, leading to false positives. QQ-plot shows the distribution of *P-*values across all ranges (from 0 to 1), providing more comprehensive information than just reporting the type 1 error rate, which only summarizes the overall distribution of *P*-values at a particular significance threshold. Importantly, the significance threshold in DGE analysis usually varies across studies due to varying numbers of tests (e.g. number of transcripts being tested) and false discovery adjustment methods. Therefore, we presented our results in QQ-plots (left panels of **Figure 2 and 3**).

**Figure 2.**
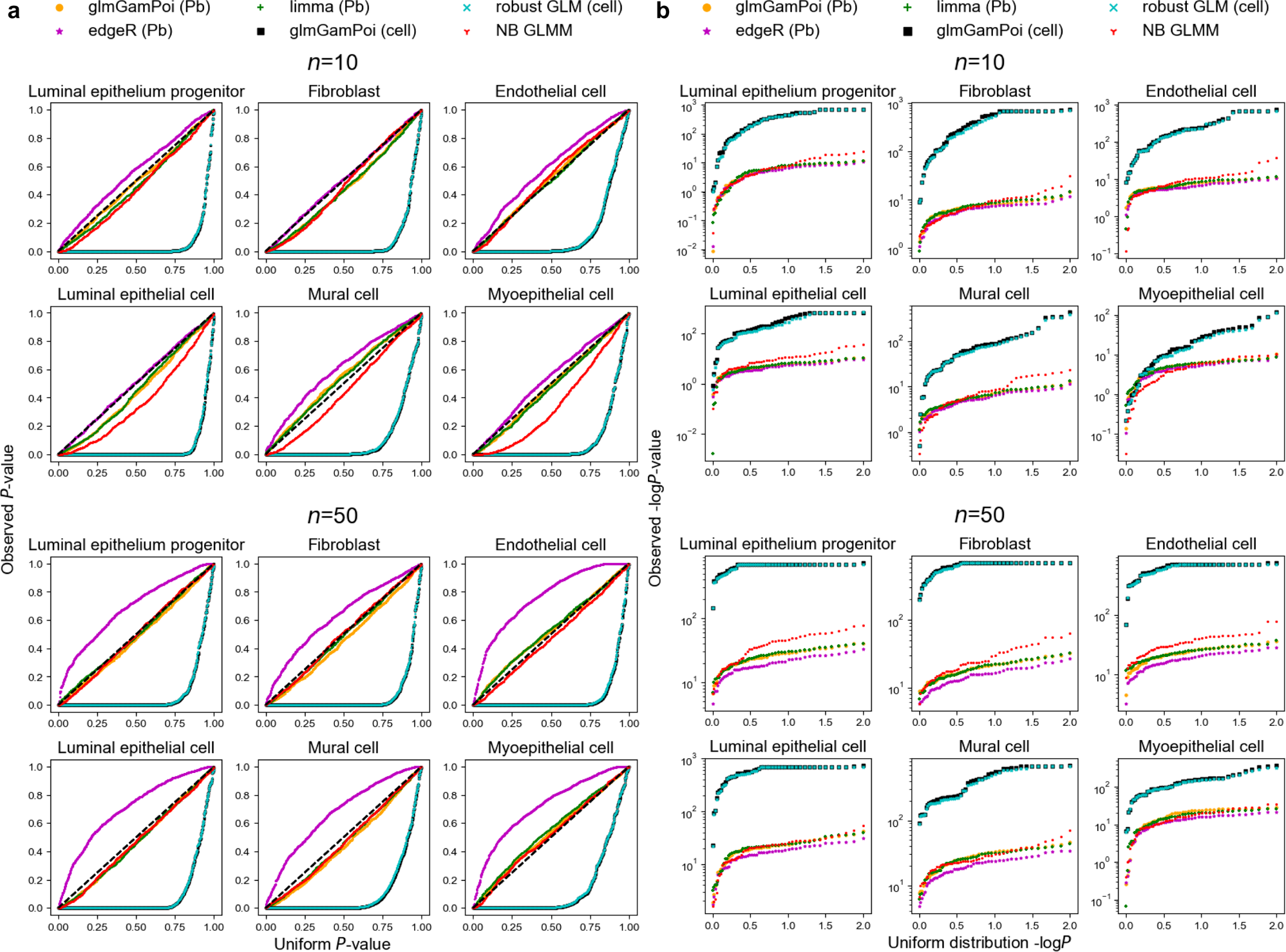
Null and power simulation when comparison group is assigned at the subject level (scenario 1) in the Reed et al. dataset. **a.** Observed versus expected *P*-values in the null simulation. **b.** Observed versus uniform -log*P* values.

**Figure 3.**
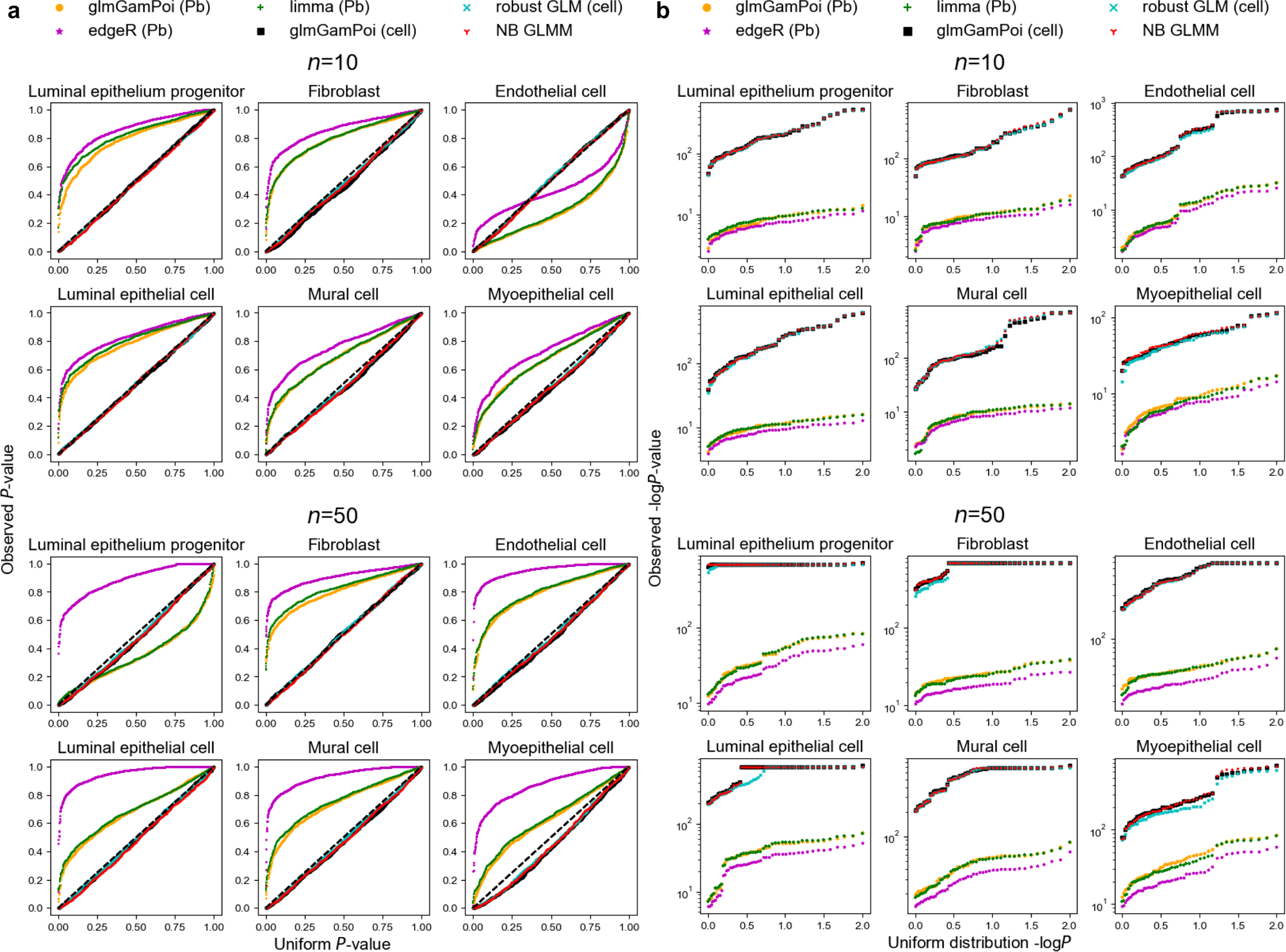
Null and power simulation when comparison group is assigned at the cell level (scenario 2) in the Reed et al. dataset. **a.** Observed versus expected *P*-values in the null simulation. **b.** Observed versus uniform -log*P* values.

In the right panel, we presented the log-transformed *P-*values generated from the power simulation. In this case, the true difference between the comparison group is set to a non-zero value, so the *P*-values are smaller than those produced from the null simulation. The more powerful the test is, the smaller the produced *P*-values. To effectively visualize the small *P*-values, we applied -log to the numbers. A more powerful test, therefore, produces a larger -log *P*-value. Although higher power offers better sensitivity, it might result from sacrificing false discovery rate. Hence, a test that has higher power without controlling type 1 error properly should be interpreted with caution (e.g. cell-wise methods in scenario 1 in the following simulations).

Overall, pseudobulk methods and NB GLMM performed well in the first scenario (**Figure 2**, **Supplementary Figure 1 and 2**). In terms of type 1 error, pseudobulk methods gave calibrated error, except edgeR gave lower error than desired in several datasets. NB GLMM’s *P*-values were deflated with small sample size (*n*=10), but they were improved with large sample size (*n*=50). In terms of power, NB GLMM was slightly more powerful than pseudobulk methods in some datasets. By contrast, cell-wise methods were severely anti-conservative, with many false positives in the null simulation. Although cell-wise methods look more powerful according to the *P-*values in the right panel of **Figure 2**, this should be interpreted along with the failure to control type 1 errors.

In the second scenario where the cell states vary within a subject, cell-wise methods and NB GLMM exhibited well-calibrated *P*-values under the null (**Figure 3**, **Supplementary Figure 3 and 4**). Pseudobulk methods, on the other hand, were generally too conservative and often produced overly large *P-*values under the null, significantly diverging from the expected distribution. In the power simulation, cell-wise methods and NB GLMM’s *P*-values were much smaller than those of pseudobulk, demonstrating these should be the methods of choice for this scenario.

### Analysis of publicly available experimental perturbation datasets

A variety of single-cell perturbation assays have been developed. As shown in the previous section, these data (scenario 2) have a different data-generating process, so DGE analysis should be performed differently from case-control single-cell studies (scenario 1). If not, the chance of missing true signal can increase substantially.

To demonstrate this point, we analyzed two perturbation datasets using the 6 methods compared in the simulations (see **Methods**). Note that these datasets have been frequently analyzed through pseudobulk approaches^10,12,16–18^. Under a stringent significance threshold (*p*<0.05/6257), pseudobulk methods rejected less than 7% transcripts, while cell-wise methods and NB GLMM rejected more than 11.5% in an interferon-g (IFN-g) stimulation experiment (**Figure 4a**). Similarly, in a CRISPR perturbation experiment, pseudobulk methods failed to find any signals, while cell-wise methods and NB GLMM found more than 10% of the genes to be differentially expressed (*p<*0.05/4807, **Figure 4b**).

**Figure 4.**
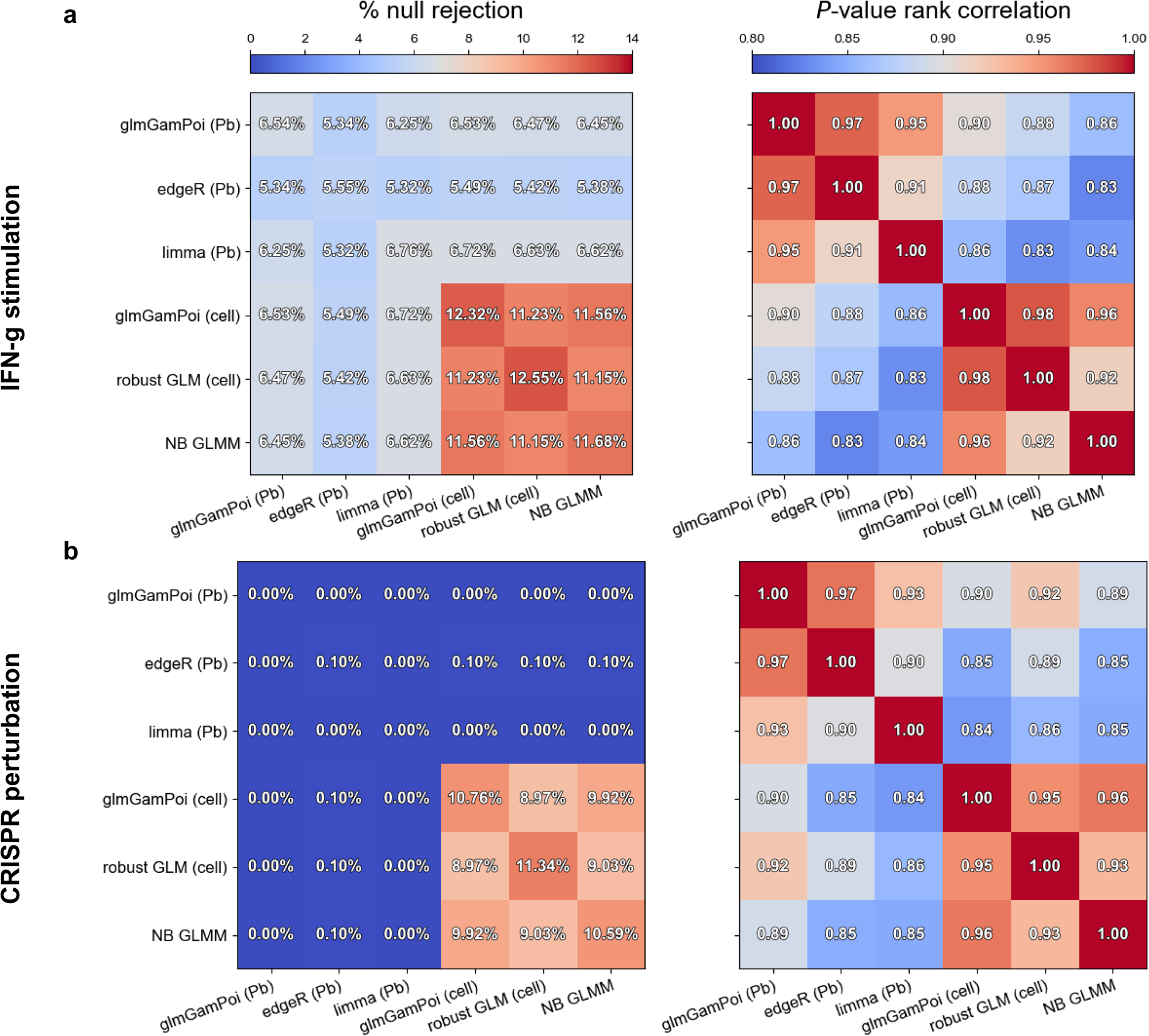
Heatmap of proportion (%) of null rejections and the Spearman correlation between methods. **a.** IFN-g stimulation of peripheral blood mononuclear cells (PBMC)^9^. **b.** CRISPR perturbation of T-cells^10^. In the left panel, the diagonal values are the proportion of rejected nulls of the methods. The non-diagonal values are the proportion of nulls that were rejected by both methods. The values of the right panel are pairwise the Spearman correlation of *P*-values.

Nevertheless, the rank of genes ordered by their *P*-values was consistent to some degree. As can be expected, the concordance was higher within pseudobulk methods and within cell-wise (or NB GLMM) methods (**Figure 4a and Figure 4b**), reaching pairwise correlations higher than 0.9 within these groups. Across these groups, the Spearman rank correlations dropped down to 0.83. Although 0.83 looks high, this imperfect (<1) correlation implies that the ranks of key genes can change.

## Discussion

We demonstrated the importance of study design in single-cell DGE analysis. The seemingly common data structure in single-cell studies, where comparison groups are discrete random variables (e.g. dummy variables in 0 and 1) and the outcome variables are the observed transcript counts, has confused researchers. The study design, the mechanism of how these discrete group variables are assigned to the cells, has a profound consequence on DGE analysis. When cell states are assigned at the subject level, cells within a subject can be treated as a single observation as a whole via pseudobulk. On the contrary, they can be treated as independent observations when each cell is assigned its state independently (**Supplementary Note**).

The ramification of ignoring study design comes in both ways. False discoveries are severely inflated when cells are treated as independent observations when they aren’t, and true DEGs are missed when independent observations are treated as aggregates. For example, pseudobulk is effective in accounting pseudoreplication bias in case-control studies but is inadequate for perturb-seq experiments. Cell-level methods are powerful and valid for perturb-seq experiments but are fragile when applied to case-control studies.

We proposed a theoretical argument on the origin of pseudoreplication bias (**Supplementary Note**). Most, if not all, statistical estimators’ variance is inversely proportional to the number of independent replicates due to the central limit theorem^19^. We’ve shown that this number is the number of samples (*n*) and the number of cells (*N*) in the first and the second scenarios, respectively. In case-control designs, therefore, the subjects are the true replicates, and the cells are fake ones, which explains the name of the pseudoreplication bias. In the second design, the estimator converges to the true parameter at a 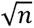-rate, so cells are true replicates in this case. Therefore, cell-wise methods correctly estimated the standard errors. Also, pseudoreplication bias does not occur in the second case.

Negative-binomial generalized linear mixed model (NB GLMM) was the only calibrated method in both designs, given a sufficient sample size. Although the high computational demand often makes it less attractive, a scalable implementation called NEBULA has been introduced^20^. One weakness is that NB GLMM often fails to converge. We found that this happens frequently in transcripts with low mean counts (<0.1) (**Supplementary Figure 5**). If NB GLMM fails to converge, replacing it with other methods for the low-expressed transcripts is a putative solution.

scRNA-seq data are generally sparse, i.e., they have many zeros^21^. Dropout has been considered a potential cause, and several methods were developed to address this phenomenon in DGE analysis^22,23^. We investigated the performance of two methods (MAST mixed model and DEsingle). In our simulations, DEsingle was anti-conservative and conservative in scenarios 1 and 2, respectively. This was expected as DEsingle infers the number of replicates directly from the dimension of the input matrix. MAST performed well when the number of subjects was small but was often conservative when the subject number grew (**Supplementary Figure 6 and 7**). Also, severe inflation of false discoveries was observed in the second scenario. More recent works suggest that the combination of the discreteness and the low mean of the count distribution causes the observed sparsity^21,24–26^. Also, zeros from the low count distribution and the dropout component may not be distinguishable solely through statistical procedures^21,24^. The latest normalization method SCTransform of Seurat also does not consider zero-inflation in UMI datasets^27^. Therefore, we raise caution on using zero-inflated models.

The cell-wise and pseudobulk methods evaluated in the paper are all distribution-robust methods. edgeR and glmGamPoi implement a quasi-likelihood ratio test ^15,28,29^. By quasi, it means that the likelihood does not require the data to exactly follow the assumed distribution (i.e. negative binomial or Poisson). Robust GLM implements a robust z-test (or equivalently, the Wald test), which has the same robustness to distributional misspecification^30–32^. limma fits a linear model, but non-Gaussian data is handled similarly, making it robust against distributional violations^33^. As single-cell data consists of diverse cell populations, a simple parametric probabilistic model may not suffice. This makes robust methods an attractive choice. Additionally, in the **Supplementary Note**, we provide a theoretical argument as to why non-dropout models still work when dropouts do occur. In **Supplementary Figure 8-10**, QQ-plots of vanilla Poisson regression and negative-binomial regression under various violations of the distributional assumptions can be found. Robust GLM is the only method that remains calibrated in all scenarios.

We expect the importance of study design to grow in future studies as the complexity of single-cell studies increases. The validity of the methods should be assessed for each design separately to prevent type 1 and 2 errors.

## Methods

### Datasets in null and power simulations

Three datasets were used: Reichart et al. (human heart)^34^, Yazar et al. (human blood)^35^, and Reed et al. (human breast)^36^. We used the cell type label provided by the author. We began with the datasets provided by the authors. For each dataset, we selected the top 6 cell types and retained donors with more than 50 cells for each cell type. This left 76, 165, and 47 donors in Reichart et al., Yazar et al., and Reed et al., respectively. No further removal of data was made. The following table contains the link to the datasets.

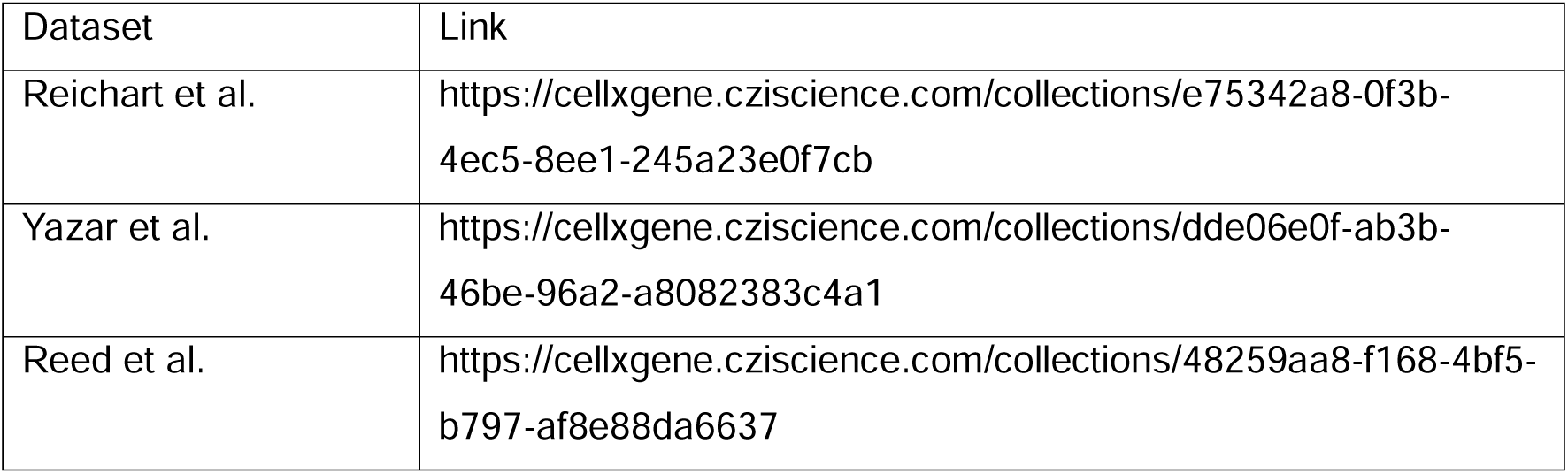

| Dataset | Link |
| --- | --- |
| Reichart et al. | https://cellxgene.cziscience.com/collections/e75342a8-0f3b-4ec5-8ee1-245a23e0f7cb |
| Yazar et al. | https://cellxgene.cziscience.com/collections/dde06e0f-ab3b-46be-96a2-a8082383c4a1 |
| Reed et al. | https://cellxgene.cziscience.com/collections/48259aa8-f168-4bf5-b797-af8e88da6637 |

The codes used in the analysis can be found at https://github.com/hanbin973/DEGpaper.

### Null simulations

In multi-subject scRNA-seq studies, cell donors are first selected from the population. Next, cells are sampled from each selected individual. In our simulation, we mimic this process by sampling individuals from the data and subsequently selecting cells from the sampled individuals. The individuals were selected with replacement to prevent artificial negative correlation between cells across individuals when the number of donors is finite^37^.

In the case of constant cell states within a donor, the first simulation assigned all cells from the same donor to the same state. Note that the state was defined as a binary variable (used R’s rbinom function). Each sampled individual was randomly allocated to either group with equal probability (p=0.5). In the second scenario, where cell state varies within a donor, cell states were assigned for each cell independently with equal probability (p=0.5, using rbinom function).

The methods evaluated in the main figure were negative binomial generalized linear mixed model (NB GLMM), glmGamPoi (with and without pseudobulk), edgeR (pseudobulk), limma (pseudobulk), and robust GLM. For NB GLMM and robust GLM, we used NEBULA and fixest software^15,20,29,33,38^, respectively. All methods were run with their default setting except for fixest (vcov=’robust’ option in fepois function). All pseudobulk was created by summing up the counts within a replicate using the pseudobulk function in glmGamPoi library. Except for limma, all methods took raw UMI counts as inputs. edgeR internally performs normalization with the raw UMI counts and voom normalization was done for limma using the ‘voom’ function of the library.

10 iterations were performed for 100 randomly selected genes in each cell-type. To prevent convergence failures in mixed models, we excluded genes with a mean smaller than 0.1 for each cell type. **Supplementary Figure 5** reports the frequency of convergence failure of NB GLMM in the Kang et al. dataset. For completeness, two implementations were tested (NEBULA and glmmTMB^39^).

### Power simulations

Power simulation was performed identically to the null simulation except for the following step. After the treatment was assigned, we randomly selected 5 genes and downsampled their count according to the prespecified fold-change (per iteration). The downsampling formula is

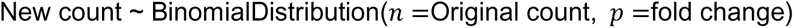

The fold change was set to 50% in all simulations.

Note that multiple genes were analyzed simultaneously per iteration to reflect the effect normalization. Methods frequently perform normalization using the total count across all transcripts present in the dataset. Earlier studies have ignored this step by testing one gene at a time ^7,14^. When a zero *P*-value occurred, we rounded them to 10^−300^ to produce the plots.

In both null and power simulations, the average number of cells of the least frequent cell types were 7481, 971 and 7511 in Reichart et al., Yazar et al., and Reed et al. datasets, respectively.

### Perturbation datasets

We adopted the code from Schmidt et al.^10^. All cells from the data were used without further filtering. Cells were divided into separate cell-types and genes with mean larger than 0.1 were selected. The links to the data and codes are listed in the following table.

| Dataset | Link |
| --- | --- |
| Kang et al. | https://www.ncbi.nlm.nih.gov/geo/query/acc.cgi?acc=GSE96583 |
| Schmidt et al. | https://zenodo.org/records/5784651 |

NB GLMM, glmGamPoi (with and without pseudobulk), edgeR (pseudobulk), limma (pseudobulk), and robust GLM (without pseudobulk) were compared. In Schmidt et al., pseudobulk was defined using the lane variable in the metadata. In Kang et al., the donor variable was used to define pseudobulk.

### Evaluation of methods considering dropouts

MAST and DEsingle were evaluated^22,23,40^. The procedure was identical to other methods in the null and the power simulations. Log normalization of the data before applying MAST was done using the scuttle library (logNormCounts function)^22,23,40^. The comparison was performed in the Yazar et al. dataset.

### Robust GLM and negative-binomial quasi-likelihood methods

Robust GLM of fixest^30^ and negative-binomial (NB) quasi-likelihood ratio test (QLRT) of edgeR and glmGamPoi^28^ are robust to distributional misspecifications. We provide a brief description of these methods. Poisson regression and negative-binomial regression are commonly implemented through maximum likelihood estimation (MLE). To perform MLE, the precise probability density (or mass) function should be known. If not, the procedure may produce bias. Although not exhaustive, bias in the point estimate (logFC in DGE) and the variance estimate of the estimator (the square of standard error) are two notable examples.

Poisson regression’s MLE estimate is correct, but the standard errors are generally deflated in overdispersed data. NB regression’s MLE estimate may suffer from both biases at the same time if data is not negative-binomial distributed. Therefore, both robust GLM and NB QLRT retain the point estimate of Poisson MLE and correct the standard error to carry out inference when the distribution is not exactly known. The correction methods are slightly different in the two cases, but the resulting *P*-values were highly concordant in our simulations (**Figure 2-3 and Supplementary Figure 1-4**). We highly recommend Wooldridge (1999)^41^ for the overview of various robust tests for count data. Lund et al. (2012)^28^ contains the details on the Bayesian shrinkage method for overdispersion estimates of NB QLRT.

Except for the name, NB QLRT and the usual NB regression are very different. NB regression estimates the regression coefficient and the overdispersion parameter simultaneously. For each iteration in optimizing the NB log-likelihood function, two parameters are optimized simultaneously. On the other hand, NB QLRT first estimates the regression coefficient using the Poisson log-likelihood function. Overdispersion is then estimated after fixing the regression coefficient obtained from the previous step^28,41,42^. The only common feature of the two methods giving NB QLRT its name is the same mean-variance relationship used to carry out inference, usually called the NB2 parameterization^43^.

## Supplementary Figure legend

**Supplementary Figure 1.**
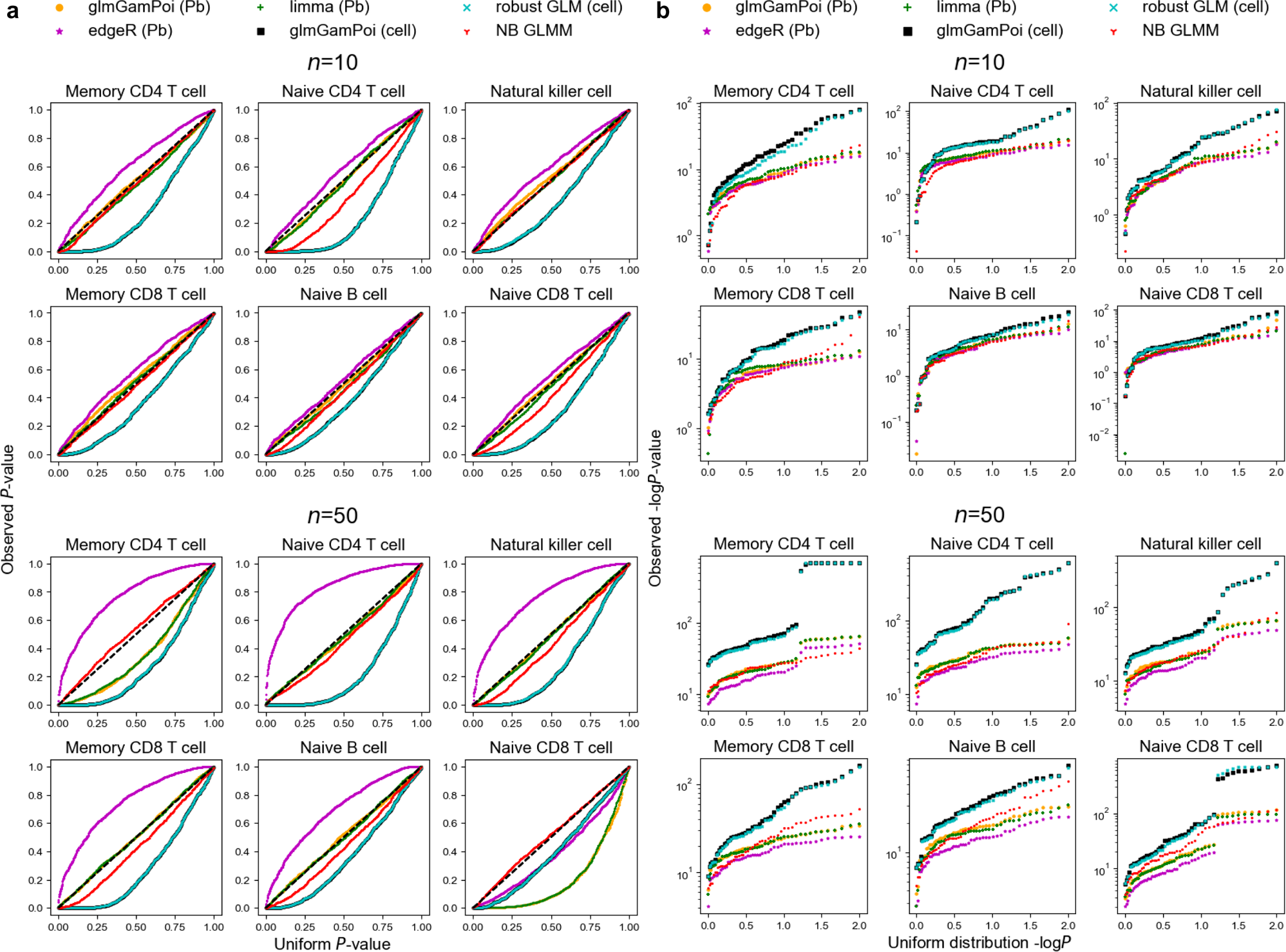
Null and power simulation when comparison group is assigned at the subject level (scenario 1) in the Yazar et al. dataset. **a.** Observed versus expected *P*-values in the null simulation. **b.** Observed versus uniform -log*P* values.

**Supplementary Figure 2.**
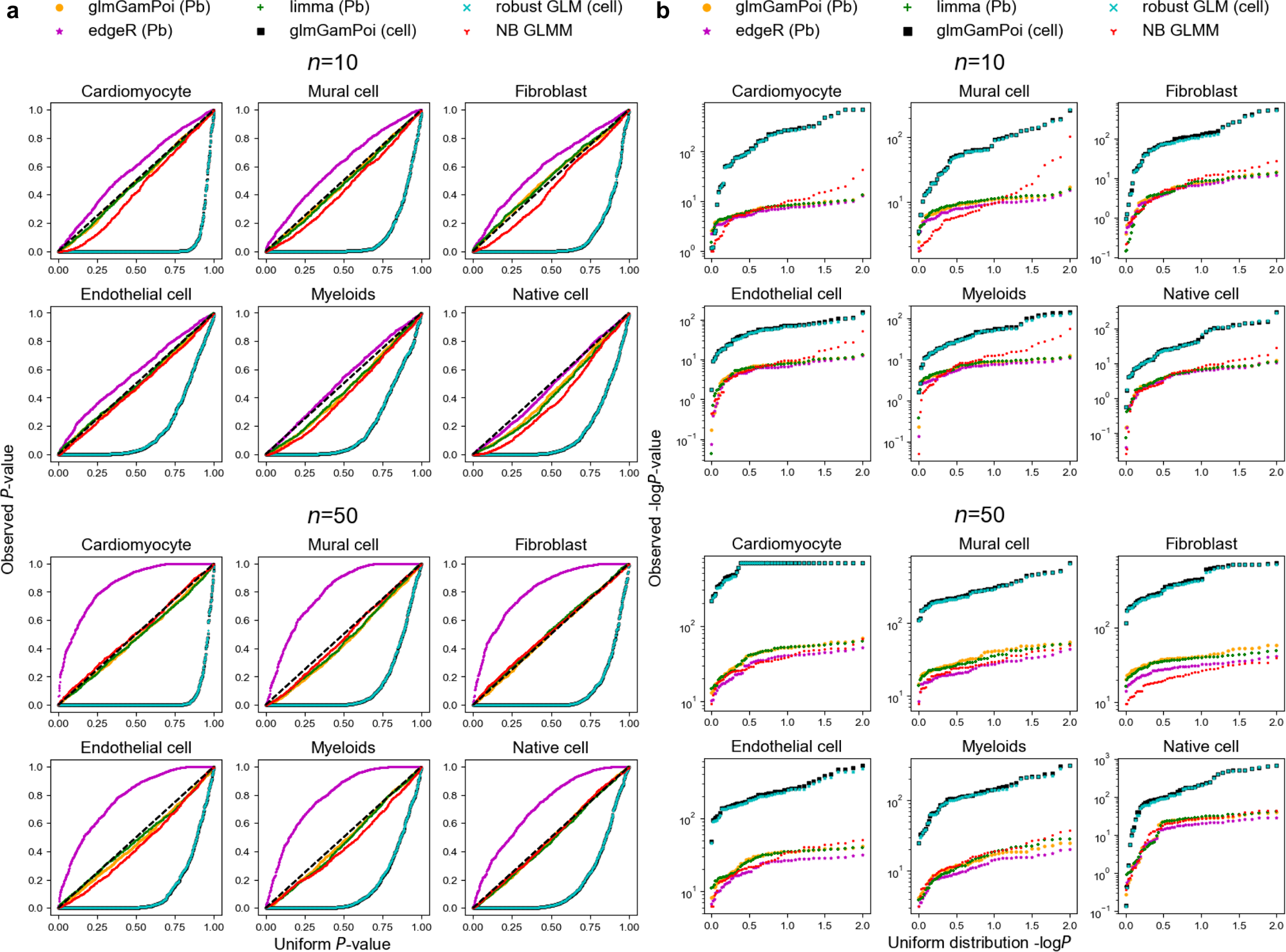
Null and power simulation when comparison group is assigned at the subject level (scenario 1) in the Reichart et al. dataset. **a.** Observed versus expected *P*-values in the null simulation. **b.** Observed versus uniform -log*P* values.

**Supplementary Figure 3.**
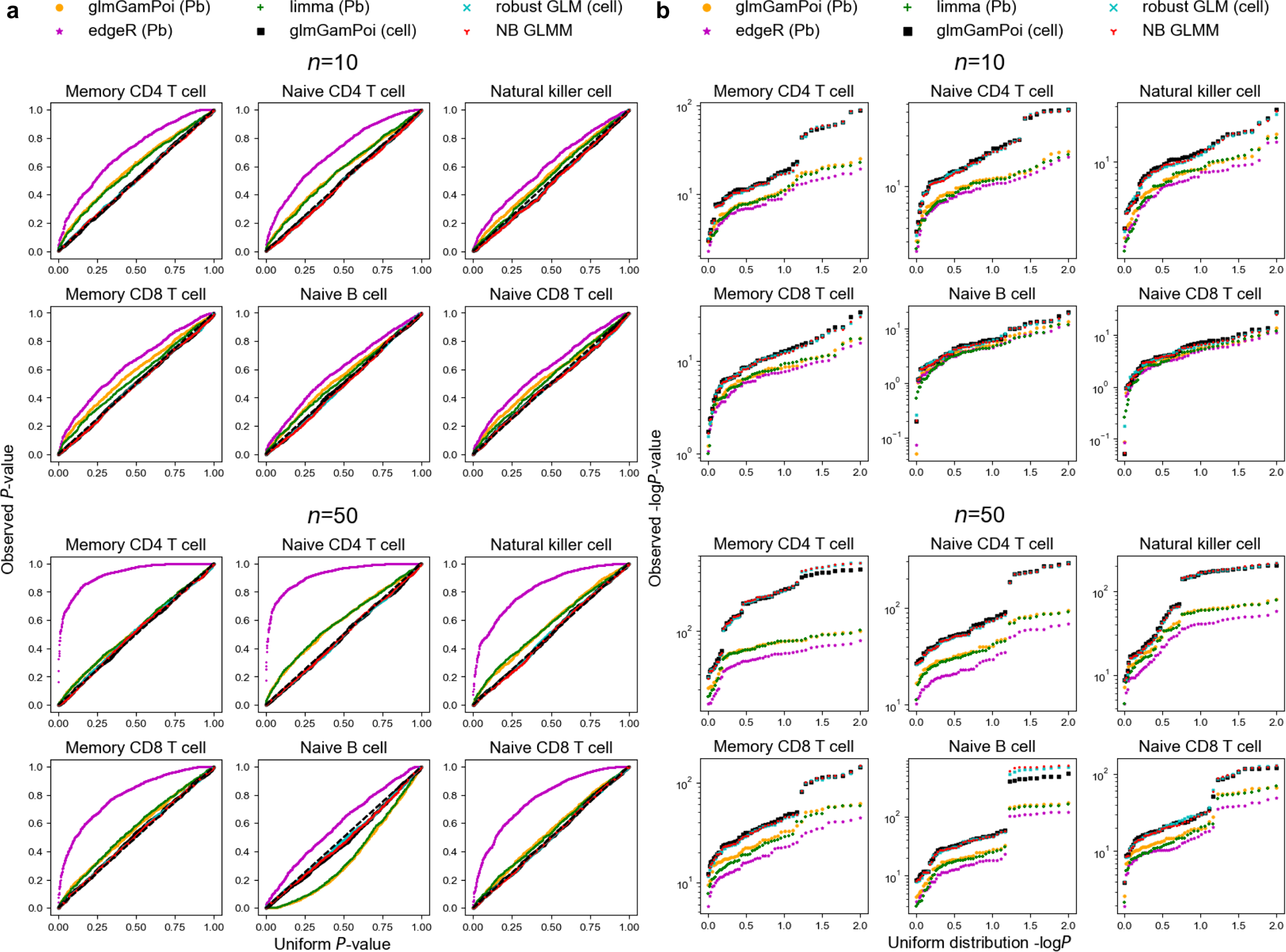
Null and power simulation when comparison group is assigned at the cell level (scenario 2) in the Yazar et al. dataset. **a.** Observed versus expected *P*-values in the null simulation. **b.** Observed versus uniform -log*P* values.

**Supplementary Figure 4.**
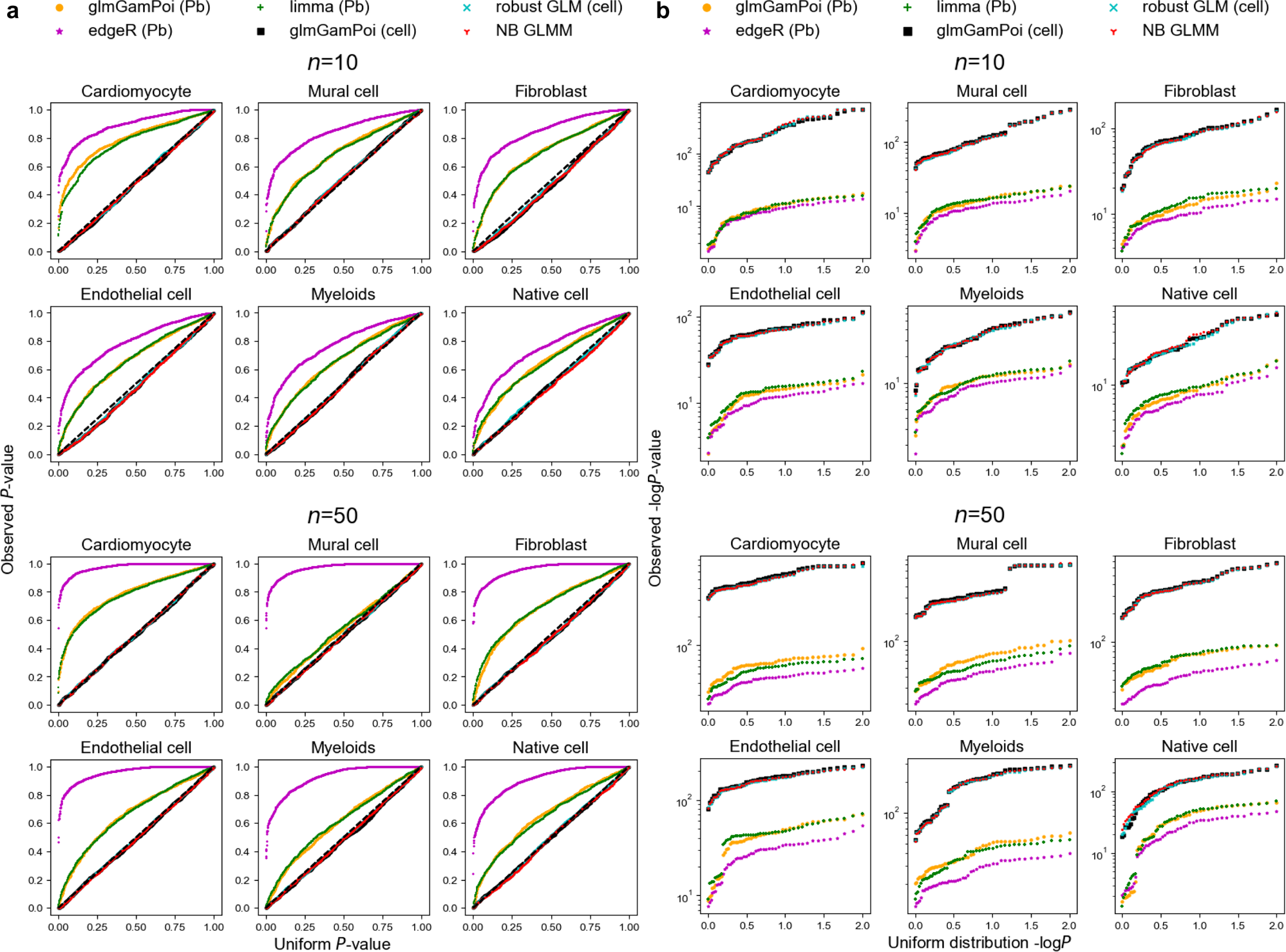
Null and power simulation when comparison group is assigned at the cell level (scenario 2) in the Reichart et al. dataset. **a.** Observed versus expected *P*-values in the null simulation. **b.** Observed versus uniform -log*P* values.

**Supplementary Figure 5.**
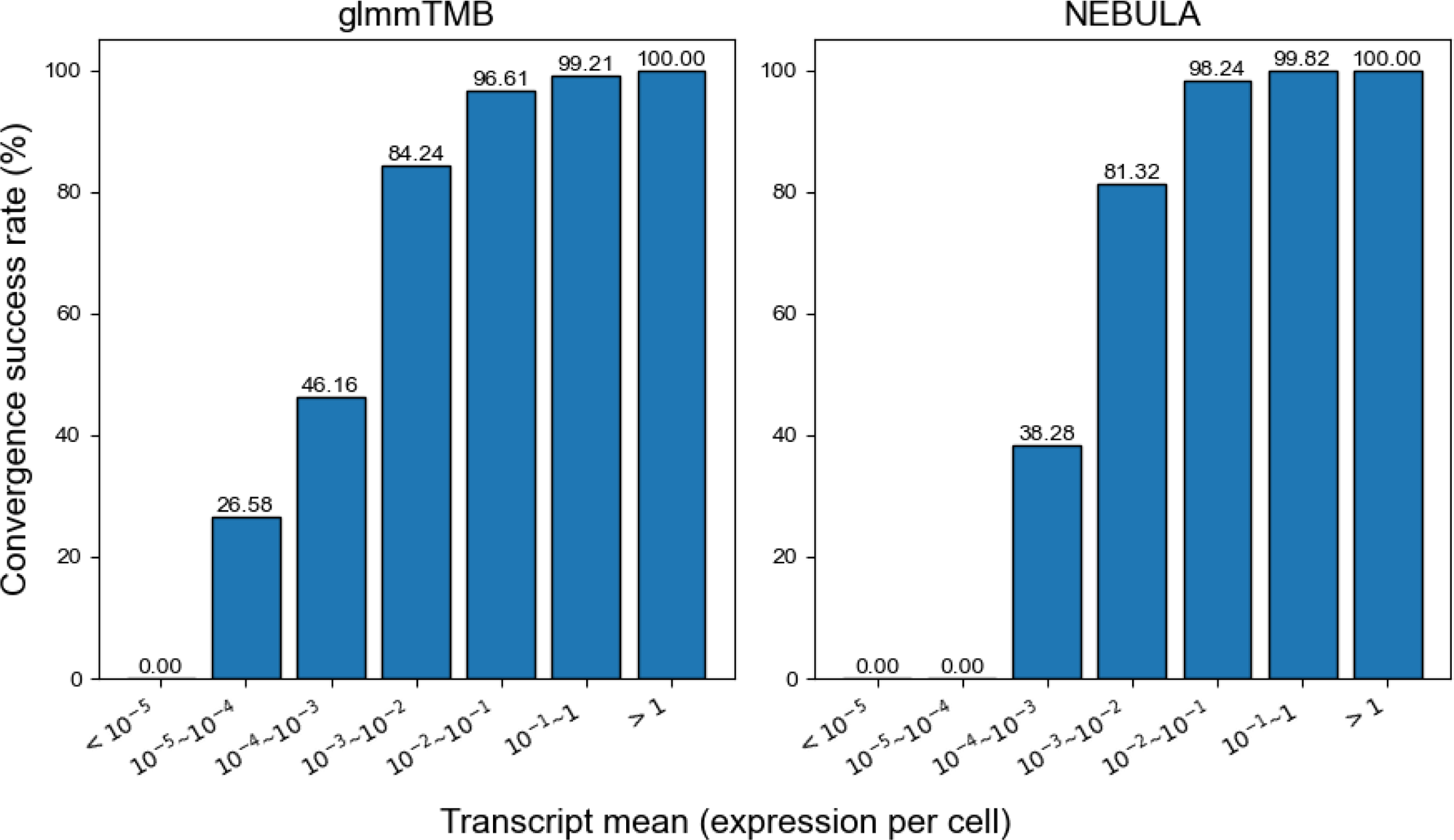
Convergence failure rate versus transcript mean of negative-binomial mixed model. **a.** glmmTMB implementation. **b.** NEBULA implementation.

**Supplementary Figure 6.**
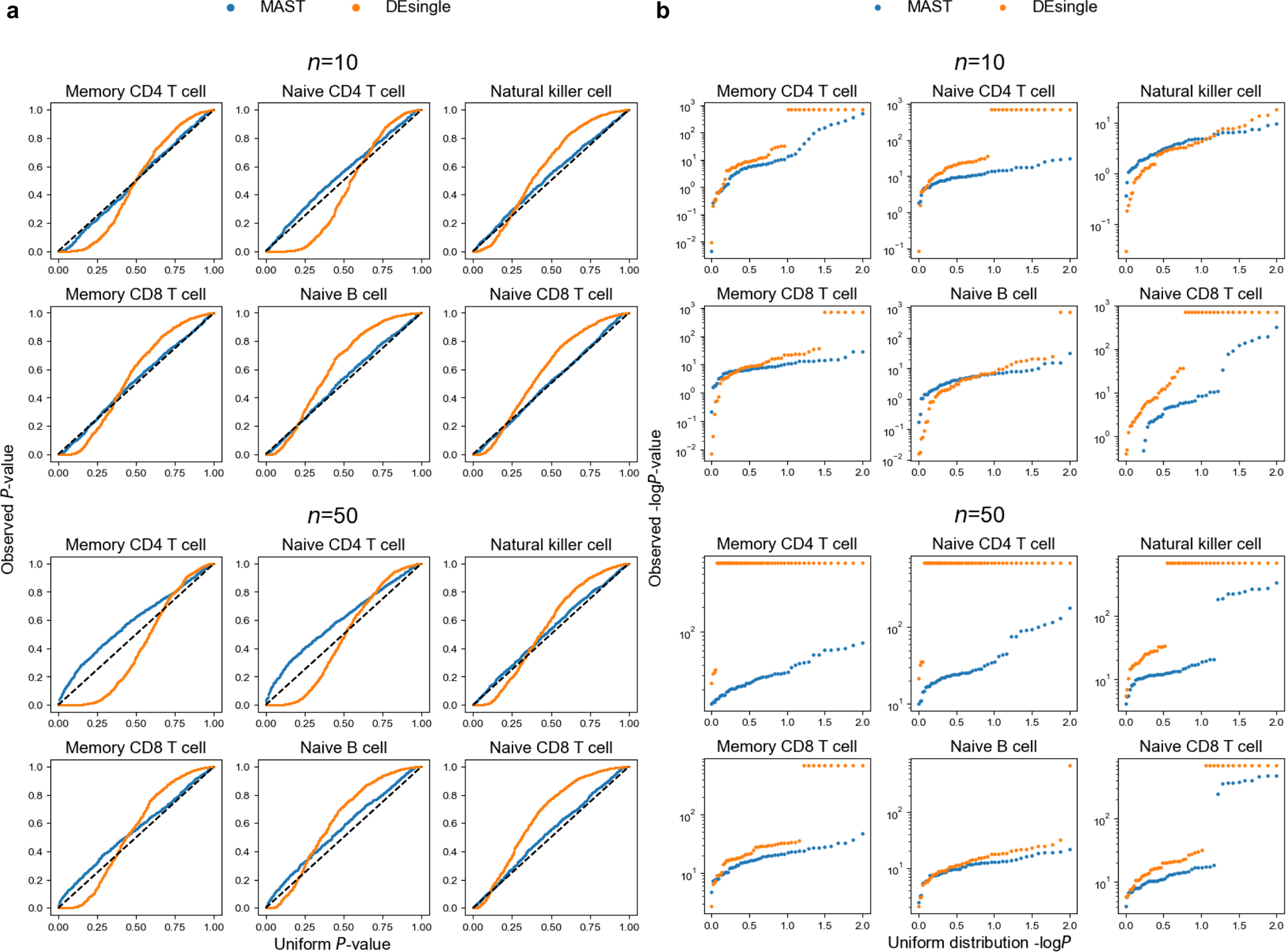
Null and power simulation when comparison group is assigned at the subject level (scenario 1) in the Yazar et al. dataset. The methods are MAST mixed-model and DEsingle. **a.** Observed versus expected *P*-values in the null simulation. **b.** Observed versus uniform -log*P* values.

**Supplementary Figure 7.**
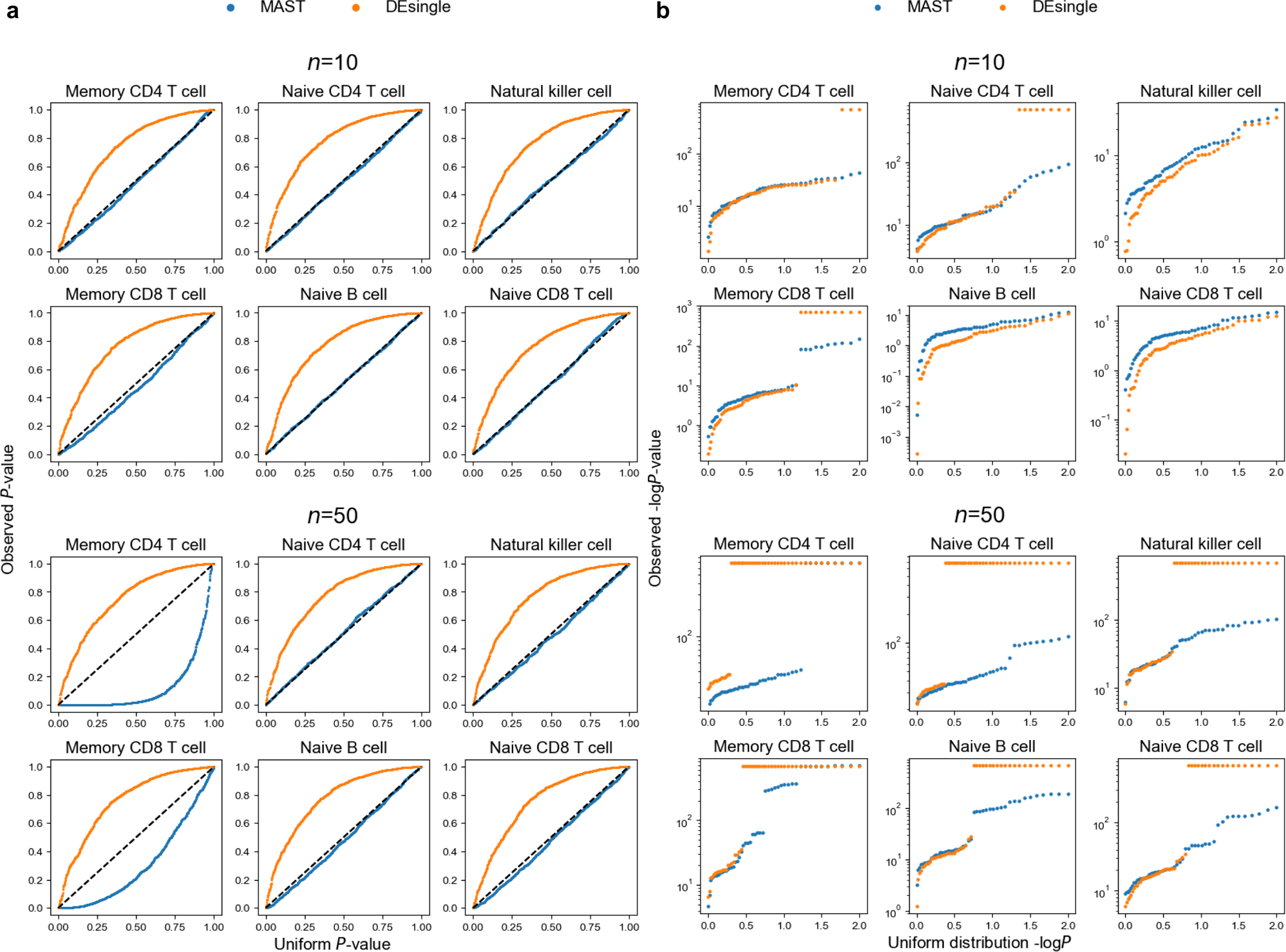
Null and power simulation when comparison group is assigned at the cell level (scenario 2) in the Yazar et al. dataset. The methods are MAST mixed-model and DEsingle. **a.** Observed versus expected *P*-values in the null simulation. **b.** Observed versus uniform -log*P* values.

**Supplementary Figure 8.**
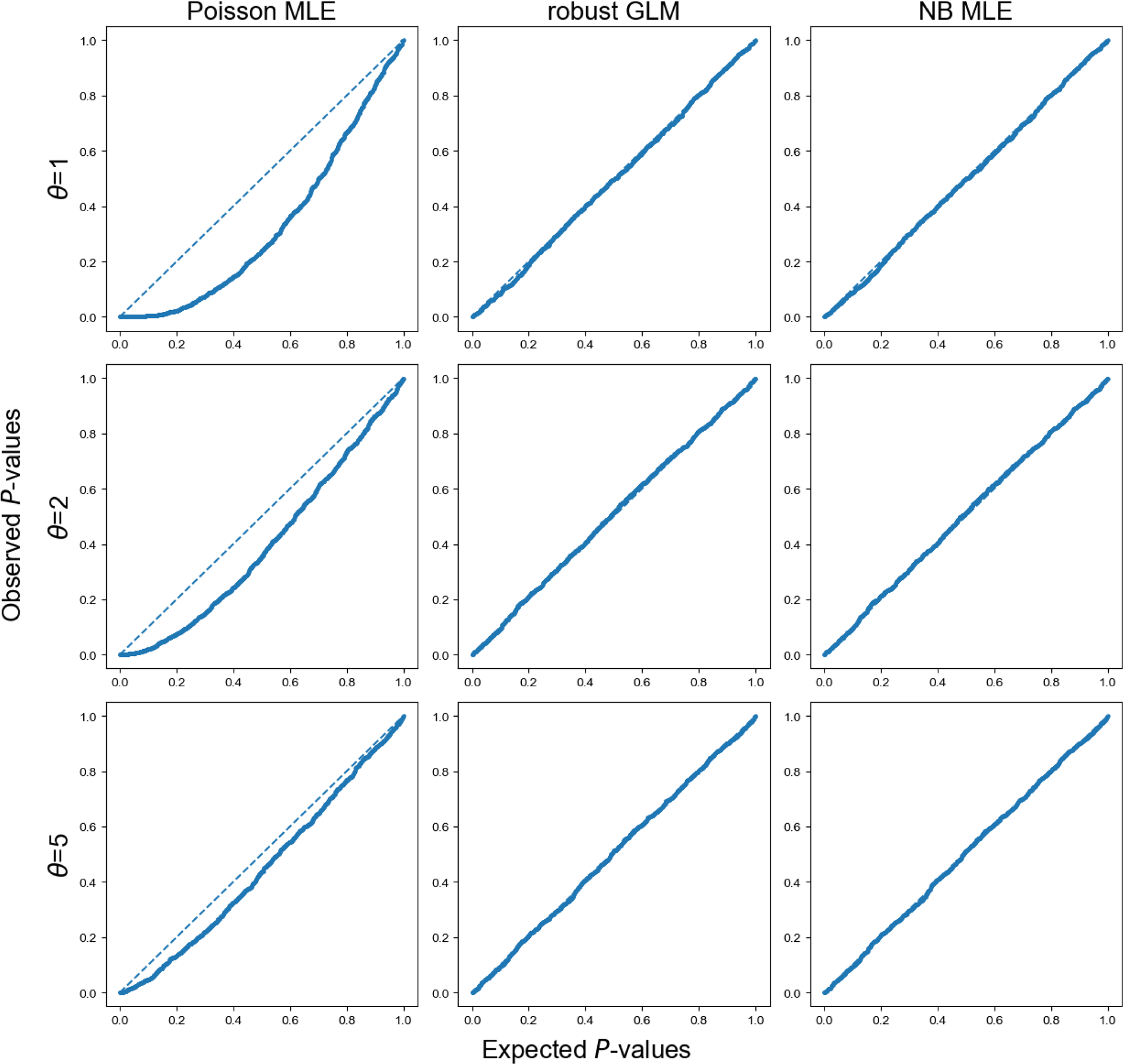
QQ-plots of Poisson MLE, robust GLM, and negative binomial (NB) MLE under negative-binomial distribution with varying dispersion parameter *θ*=1,2,5 in *Var*(*y*) = *µ* + *µ*^2^/*θ* (NB2 parameterization).

**Supplementary Figure 9.**
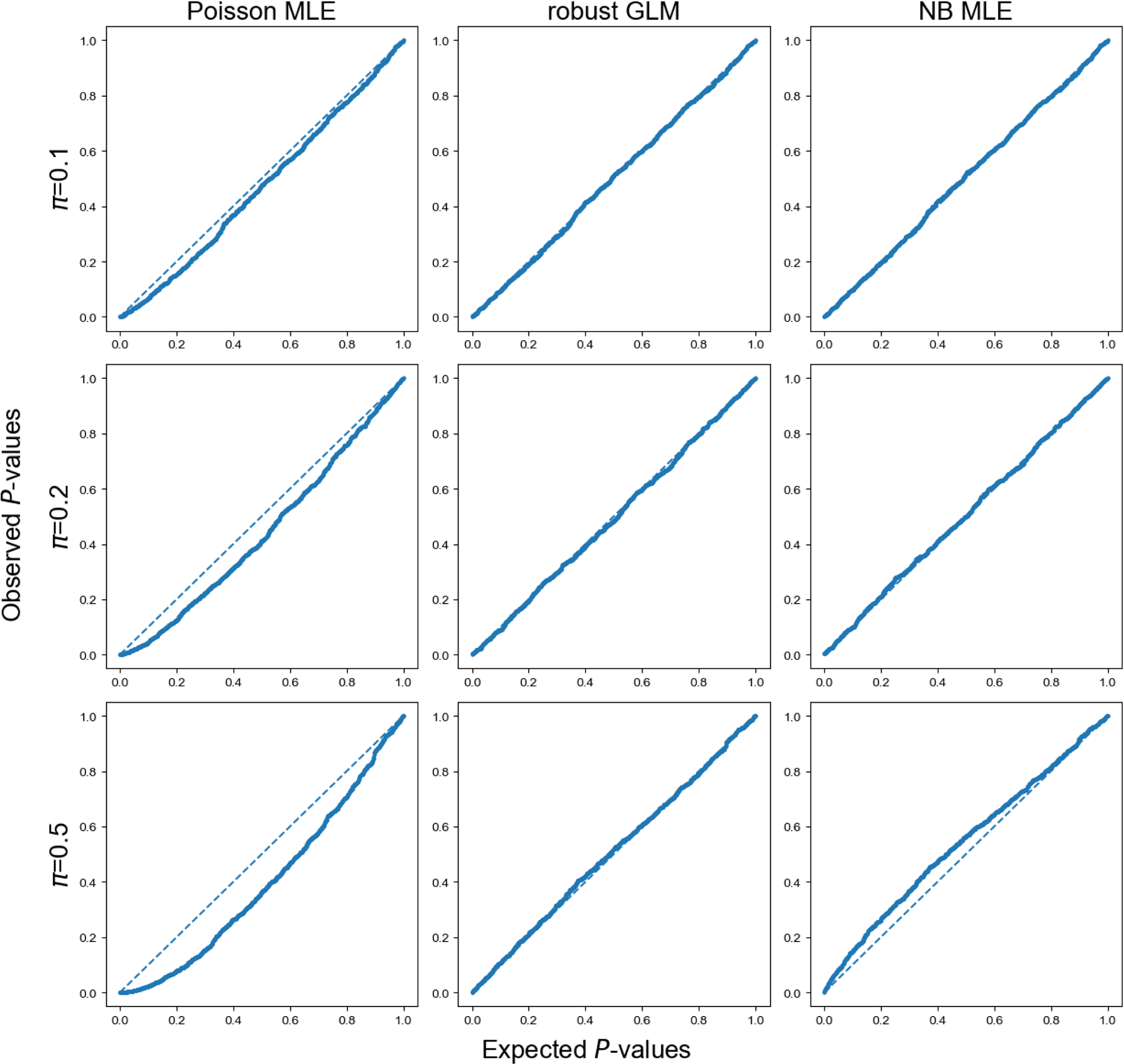
QQ-plots of Poisson MLE, robust GLM, and negative binomial (NB) MLE under zero-inflated Poisson distribution with varying zero inflation rate *π*=0.1, 0.2, 0.5.

**Supplementary Figure 10.**
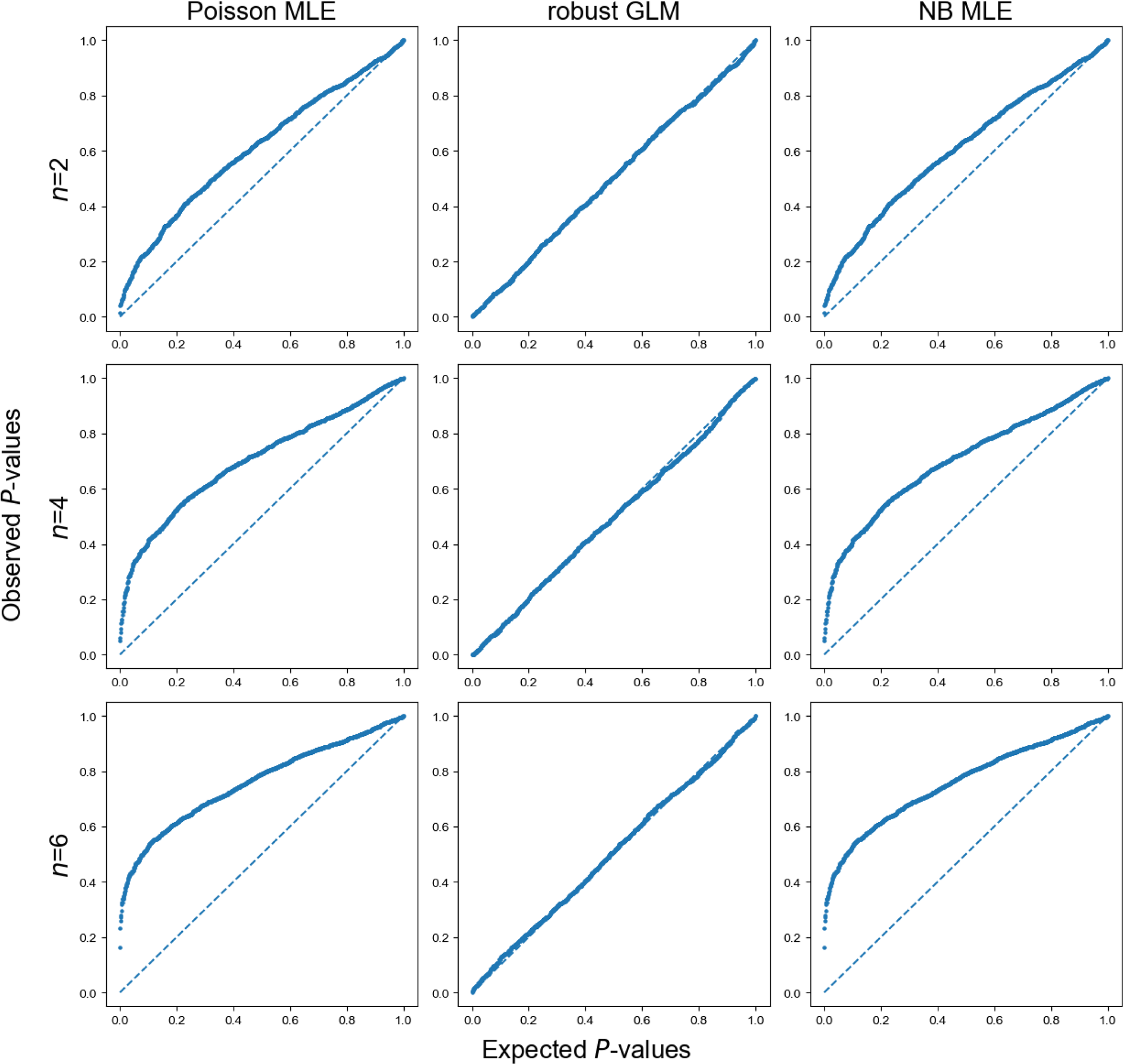
QQ-plots of Poisson MLE, robust GLM, and negative binomial (NB) MLE under a mixture of Poisson distributions with varying numbers of mixture components *n*=2,4,6.

## Notes

### Competing Interest Statement

Buhm Han is the CTO of the Genealogy Inc.

### Summary of Updates

Main text update

