## Supplementary File for "General solution to inflated type I and II errors in multi-subject single-cell differential gene expression analysis"

### Supplementary Note

Hanbin Lee

November 20, 2023

#### 1 Setup

We assume that the expression counts obey the following equation.

$$g\left(\mathbb{E}\left[Y_{ijk} \mid X_{ij}, \alpha_{ik}\right]\right) = \beta_{0k} + X_{ij}\beta_{1k} + \alpha_{ik} \tag{1}$$

$i$ is the index of subjects, $j$ is the index of cells, and $k$ is the index of genes. $Y_{ijk}$ is the expression count, $X_{ij}$ is the cell status (0 or 1 for the binary state to compare), and $\alpha_{ik}$ is the subject-specific effect on expression. $g$ is the link function. Alternatively, the equation can be written as

$$Y_{ijk} = g^{-1}\left(\beta_{0k} + X_{ij}\beta_{1k} + \alpha_{ik}\right) + \epsilon_{ijk} \tag{2}$$

where $\epsilon_{ijk} = Y_{ijk} - \mathbb{E}\left[Y_{ijk} \mid X_{ij}, \alpha_{ik}\right]$. Note that $\mathbb{E}\left[\epsilon_{ijk} \mid X_{ij}\right] = 0$.

The link function $g$ is defined based on the assumed model. Linear models preceded by log-like normalization (e.g., limma) follow the above equation with $g(x) = x$, the identity function. Poisson regression, negative binomial regression, and their associated mixed models use $g(x) = \log x$, the log function. For the sake of generality, the following derivations do not assume a particular probability distribution.

When $g(x) = \log x$, $\beta_{1k}$ is the log fold-change (logFC) which is the parameter of interest. Even for the $g(x) = x$ case, after suitable log transformation of $Y_{ijk}$, $\beta_{1k}$ is an approximation of the logFC. In the following sections, the proofs assume $g(x) = \log x$ because the derivation is more difficult than $g(x) = x$. The proofs for the $g(x) = x$ case can be obtained from the $g(x) = \log x$ case after removing all exp and log functions.

#### 2 Autocorrelation and pseudoreplication

When the expressions in the cells within a subject are correlated (not independent), we call it *autocorrelation.* In equation (1), $\alpha_{ik}$ is the term responsible for the autocorrelation. As $\alpha_{ik}$ varies across subjects, and subjects are randomly selected from the population, $\alpha_{ik}$ is a random variable with non-zero variance. If it were constant, the intercept $\beta_{0k}$ absorbs the term leaving no autocorrelation. Therefore, autocorrelation can only exist when there are two or more subjects since all cells will have the same $\alpha_{ik}$ when there is only one subject. We assume that autocorrelation exists in the data (otherwise, everything simplifies, and applying the cell-wise method to any data should work).

In literature, autocorrelation is attributed as the cause of *pseudoreplication bias* [3, 2]. There are several observations related to the bias. First, the data should have of multiple subjects so that autocorrelation is present. Second, when a cell-wise method is applied to a multi-subject data, the false discovery rate (FDR) increases substantially. Third, aggregating the cells into a pseudobulk ameliorates the inflated FDR. Another alternative is to use a mixed model. It follows that autocorrelation causes the bias, and the two methods resolve it by taking autocorrelation into account. Following the same logic, the failure of cell-wise methods comes from ignoring autocorrelation.

Autocorrelation explains the name of the bias. *Replicate* refers to the independent units of observation from a study [1, 3]. As cells are mutually correlated, they are not replicates. Instead, they are called *pseudoreplicates* for not being independent. Since cell-wise methods treat cells (which are pseudoreplicates) as independent observations (i.e. replicates), it makes sense to call the high FDR of cell-wise methods pseudoreplication bias because it stems from mistaking pseudoreplicates as independent replicates.

The argument gives the impression that pseudoreplication bias happens whenever cell-wise method is used in the presence of autocorrelation. However, we have shown that cell-wise methods are well-calibrated in the second scenario discussed in the main text. To close the gap between the previous understanding of pseudoreplication bias and our observations, we suggest a more narrow and precise definition of pseudoreplication bias based on theoretical speculation.

### 3 Pseudoreplication bias revisited

We show below that the pseudoreplication bias occurs only if the three conditions are all met: (1) multi-subject design (with autocorrelation), (2) constant cell states within a subject (namely scenario 1), and (3) cell-wise method used. We recast the bias in the language of estimator variance to analytically prove our argument.

An estimator $\widehat{\beta}_{1k}$ guesses the parameter $\beta_{1k}$ with a finite amount of data. Since the estimator is a function of the data which is a random variable, it is also random. It means that the estimator varies to a certain degree for each experiment. An estimator's variance measures the variability of the estimator under repeated (but identical) experiments. Small variance means less variability, i.e. the value of the estimator is likely to concentrate in a smaller range under repeated experiments. However, we usually have a single realization of potentially many experiments so the true variance is unknown. The estimated variance is then used to compute the $P$-value to assess statistical significance. In short, there is the true variance of an estimator under repeated experiments, $\mathrm{Var}\left(\widehat{\beta}_{1k}\right)$, and the estimated variance $\widehat{\mathrm{Var}}\left(\widehat{\beta}_{1k}\right)$ that is used in practice because the true variance is only theoretically known.

A method to detect DEG is equipped with an estimator $\widehat{\beta}_{1k}$ and its corresponding variance estimate $\widehat{\mathrm{Var}}\left(\widehat{\beta}_{1k}\right)$. If the estimate $\widehat{\mathrm{Var}}\left(\widehat{\beta}_{1k}\right)$ is a proper guess for $\mathrm{Var}\left(\widehat{\beta}_{1k}\right)$, the test is valid and produces calibrated $P$-values. Conversely, if the method underestimates the true variance, it produces deflated $P$-values, leading to an inflation of false discoveries. For example, $z$-test computes

$$Z = \frac{\widehat{\beta}_{1k}}{\widehat{\mathrm{Var}}\left(\widehat{\beta}_{1k}\right)} \tag{3}$$

and tests if $|Z|$ exceeds a pre-specified threshold to declare significance. Thus, an underestimation of $\widehat{\mathrm{Var}}\left(\widehat{\beta}_{1k}\right)$ increases $|Z|$ leading to more false discoveries even if $\widehat{\beta}_{1k}$ is unbiased. Other tests like $\chi^2$-test, Wald test, and likelihood ratio test (LRT) also utilize the similar variance estimate. Thus, the significance is exaggerated whenever the variance is underestimated, regardless of the test method used.

For any cell-wise method, the estimated variance of its estimator is always

$$\widehat{\mathrm{Var}}\left(\widehat{\beta}_{1k}\right) = \mathcal{O}\left(\frac{1}{N}\right) \tag{4}$$

where $N$ is the number of cells, which is a consequence of assuming that $N$ cells are independent observations (replicates). Unfortunately, the true variance has different scales depending on the study design. Thus, the scale of the true variance and the estimated variance may not coincide.

**Theorem 1.**

$$\operatorname{Var}\left(\widehat{\beta}_{1k}\right) = \begin{cases} \mathcal{O}\left(\frac{1}{n}\right) & : X_{ij} \textit{ is constant within } i\textit{, Scenario 1} \\ \mathcal{O}\left(\frac{1}{N}\right) & : X_{ij} \textit{ varies within } i\textit{, Scenario 2} \end{cases} \tag{5}$$

*for any unbiased estimator* $\widehat{\beta}_{1k}$ *for the true logFC* ($\beta_{1k}$)*.* $n$ *and* $N$ *are the number of subjects and cells, respectively.*

The theorem shows that the variance estimate of any cell-wise method (4) is a severe underestimate of the true variance under scenario 1, which should increase the chance of false discovery to a large extent. This matches our previous understanding of pseudoreplication bias but with more restrictive conditions: FDR increases when cell-wise method is used in a multi-subject study but only in the first scenario where the underestimation of variance happens.

**Corollary 1.** *Pseudoreplication bias is the inflation of FDR produced by cell-wise methods in scenario 1. The inflated FDR comes from the underestimation of the estimator's variance. The following equation gives the degree of underestimation.*

$$\widehat{\operatorname{Var}}\left(\widehat{\beta}_{1k}\right) = \mathcal{O}\left(\frac{1}{N}\right) \ll \mathcal{O}\left(\frac{1}{n}\right) = \operatorname{Var}\left(\widehat{\beta}_{1k}\right) \tag{6}$$

*since* $n \ll N$*.*

**Theorem 1** also explains why pseudobulk methods are an adequate solution to the inflated FDR in scenario 1. Pseudobulk methods treat subjects as independent replicates. Therefore, the estimated variance of the estimator has the following scale

$$\widehat{\operatorname{Var}}\left(\widehat{\beta}_{1k}\right) = \mathcal{O}\left(\frac{1}{n}\right) \tag{7}$$

which matches the scale of the true variance of $\widehat{\beta}_{1k}$.

Finally, the calibrated performance of cell-wise methods in the second scenario can be understood. The scale of cell-wise methods in equation (4) equals that of the true variance in the second scenario $\mathcal{O}(1/N)$. This implies that cells are true replicates in the second scenario. Therefore, autocorrelation alone is not enough to determine the true independent unit of a study.

**Corollary 2.** *The scale of the estimator variance of cell-wise methods equals to the scale of the true variance under scenario 2.*

$$\widehat{\text{Var}}\left(\widehat{\beta}_{1k}\right) = \mathcal{O}\left(\frac{1}{N}\right) = \text{Var}\left(\widehat{\beta}_{1k}\right) \tag{8}$$

*Note that the equality sign '=' here means they coincide in terms of scale, not being numerically the same.*

Now we move on to the proofs of **Theorem 1**.

*Proof.* The proof of the claim (**Theorem 1**) comes in two steps. First, we analyze the behavior of $\widehat{\beta}_{1k}$ conditioned on the sampled subjects. This fixes $\boldsymbol{\alpha}_k$, the collection of all $\alpha_{ik}$, removing the autocorrelation between the cells. Next, we let $\boldsymbol{\alpha}_k$ vary and observe the full behavior. The law of iterated variance gives

$$\text{Var}\left(\widehat{\beta}_{N,1k}\right) = \text{Var}\left(\mathbb{E}\left[\widehat{\beta}_{N,1k} \mid \mathbf{X}, \boldsymbol{\alpha}_k\right]\right) + \mathbb{E}\left[\text{Var}\left(\widehat{\beta}_{N,1k} \mid \mathbf{X}, \boldsymbol{\alpha}_k\right)\right] \tag{9}$$

where $\mathbf{X}$ is the collection of all $X_{ij}$ (cell status of cell $j$ of individual $i$). The subscript $N$ means the estimator is computed from $N$ cells. Conditioning on $\mathbf{X}$ and $\boldsymbol{\alpha}_k$ gives the effect of fixing the sampled subjects. As explained earlier, we analyze $\mathbb{E}\left[\widehat{\beta}_{N,1k} \mid \mathbf{X}, \boldsymbol{\alpha}_k\right]$ and $\text{Var}\left(\widehat{\beta}_{N,1k} \mid \mathbf{X}, \boldsymbol{\alpha}_k\right)$ first which describe the behavior of the estimator with $\mathbf{X}$ and $\boldsymbol{\alpha}_k$ fixed.

**Proposition 1.** *The behavior of the second term,* $\text{Var}\left(\widehat{\beta}_{N,1k} \mid \mathbf{X}, \boldsymbol{\alpha}_k\right)$*, is design-independent, and*

$$\text{Var}\left(\widehat{\beta}_{N,1k} \mid \mathbf{X}, \boldsymbol{\alpha}_k\right) = \mathcal{O}\left(\frac{1}{N}\right) \tag{10}$$

*Proof.* As $\mathbf{X}$ and $\boldsymbol{\alpha}_k$ are fixed, the cells are independent replications. This is because the only remaining variation of $\widehat{\beta}_{N,1k}$ after fixing $\mathbf{X}$ and $\boldsymbol{\alpha}_k$ comes from the independent sampling of cells within each subject. Then the central limit theorem of independent observation of cells gives the desired result as $N \to \infty$.

⊠

Subsequently,

$$\mathbb{E}\left[\mathrm{Var}\left(\widehat{\beta}_{N,1k} \mid \mathbf{X}, \boldsymbol{\alpha}_k\right)\right] = \mathcal{O}\left(\frac{1}{N}\right) \tag{11}$$

since the expectation of an $\mathcal{O}(1/N)$ variable is $\mathcal{O}(1/N)$.

The interesting part that exhibits design-dependent behavior is the first term, Var $\left(\mathbb{E}\left[\widehat{\beta}_{N,1k} \mid \mathbf{X}, \boldsymbol{\alpha}_k\right]\right)$. The conditional expectation, $\mathbb{E}\left[\widehat{\beta}_{N,1k} \mid \mathbf{X}, \boldsymbol{\alpha}_k\right]$ is the expected value of $\widehat{\beta}_{N,1k}$ provided the subjects are fixed. It is difficult to directly expand this quantity. However, it is doable if we consider the conditional mean expression of a specific cell. Under the condition with $\mathbf{X}$ and $\boldsymbol{\alpha}_k$ fixed, assume that we randomly select one cell (cell $j'$ of subject $i'$). If we consider the expected expression of gene $k$ of this cell:

$$\begin{aligned}
&\mathbb{E}\left[Y_{i'j'k} \mid X_{i'j'}, \mathbf{X}, \boldsymbol{\alpha}_k\right] \\
&= \mathbb{E}\left[\mathbb{E}\left[Y_{i'j'k} \mid X_{i'j'}, \alpha_{i'k}, \mathbf{X}, \boldsymbol{\alpha}_k\right] \mid X_{i'j'}, \mathbf{X}, \boldsymbol{\alpha}_k\right] \quad \text{(law of total expectation)} \\
&= \mathbb{E}\left[\exp\left(\beta_{0k} + X_{i'j'}\beta_{1k} + \alpha_{i'k}\right) \mid X_{i'j'}, \mathbf{X}, \boldsymbol{\alpha}_k\right] \quad \text{(by the definition of } Y_{i'j'k} \text{ in eq. (1))} \\
&= \exp\left(\beta_{0k} + X_{i'j'}\beta_{1k}\right) \cdot \mathbb{E}\left[\exp\left(\alpha_{i'k}\right) \mid X_{i'j'}, \mathbf{X}, \boldsymbol{\alpha}_k\right] \quad \text{(constants pulled out)} \\
&= \exp\left(\beta_{0k} + X_{i'j'}\beta_{1k} + \log \mathbb{E}\left[\exp\left(\alpha_{i'k}\right) \mid X_{i'j'}, \mathbf{X}, \boldsymbol{\alpha}_k\right]\right)
\end{aligned} \tag{12}$$

We further analyze $\mathbb{E}\left[\exp\left(\alpha_{i'k}\right) \mid X_{i'j'}, \mathbf{X}, \boldsymbol{\alpha}_k\right]$. This is the design-dependent term of interest. Let $S_{i'j'} = 1, \ldots, n$ be the subject membership of cell $j'$. (That is, $S_{i'j'} = i'$. However the proof will look easier to read with this.) $\mathbb{I}(\,\cdot\,)$ is the indicator function.

$$\begin{aligned}
&\mathbb{E}\left[\exp\left(\alpha_{i'k}\right) \mid X_{i'j'}, \mathbf{X}, \boldsymbol{\alpha}_k\right] \\
&= \mathbb{E}\left[\mathbb{E}\left[\exp\left(\alpha_{i'k}\right) \mid S_{i'j'}, X_{i'j'}, \mathbf{X}, \boldsymbol{\alpha}_k\right] \mid X_{i'j'}, \mathbf{X}, \boldsymbol{\alpha}_k\right] \quad \text{(law of total expectation)} \\
&= \mathbb{E}\left[\mathbb{E}\left[\exp\left(\alpha_{i'k}\right) \mid S_{i'j'}, \mathbf{X}, \boldsymbol{\alpha}_k\right] \mid X_{i'j'}, \mathbf{X}, \boldsymbol{\alpha}_k\right] \quad (X_{i'j'} \perp \exp(\alpha_{i'k}) \mid S_{i'j'}) \\
&= \mathbb{E}\left[\sum_{i=1}^{n} \exp(\alpha_{ik}) \mathbb{I}(S_{i'j'} = i) \,\middle|\, X_{i'j'}, \mathbf{X}, \boldsymbol{\alpha}_k\right] \quad (\alpha_{ik} \text{ is constant given } S_{i'j'}) \\
&= \sum_{i=1}^{n} \exp(\alpha_{ik}) \cdot \mathbb{E}\left[\mathbb{I}(S_{i'j'} = i) \mid X_{i'j'}, \mathbf{X}, \boldsymbol{\alpha}_k\right] \\
&= \sum_{i=1}^{n} \exp(\alpha_{ik}) \cdot \mathbb{P}(S_{i'j'} = i \mid X_{i'j'}, \mathbf{X}, \boldsymbol{\alpha}_k) \quad \text{(expectation of an indicator is probability)}
\end{aligned} \tag{13}$$

In scenario 1, $X_{i'j'} = X_{i'}$ because the cells from subject $i'$ all have the same value. Given $X_{i'} = 1$, $\alpha_{i'k}$ is sampled from $\boldsymbol{\alpha}_k$ such that $X_i = 1$, and vice versa. Therefore,

$$\mathbb{P}(S_{i'j'} = i \mid X_{i'} = x, \mathbf{X}, \boldsymbol{\alpha}_k) = \begin{cases} N_i/(\sum_{h:X_h=x} N_h) & : X_i = x \\ 0 & : X_i \neq x \end{cases} \tag{14}$$

which is the probability among subjects $X_i = x$ or zero, otherwise. $N_i$ is the number of cells in subject $i$, such that $N = N_1 + ... + N_n$.

In scenario 2, the fact that $X_{i'j'}$ varies independently within a subject implies $S_{i'j'} \perp X_{i'j'} \mid \mathbf{X}, \boldsymbol{\alpha}$. Hence,

$$\mathbb{P}(S_{i'j'} = i \mid X_{i'j'} = x, \mathbf{X}, \boldsymbol{\alpha}_k) = \mathbb{P}(S_{i'j'} = i \mid \mathbf{X}, \boldsymbol{\alpha}_k) = \frac{N_i}{\sum_{h=1}^n N_h} \tag{15}$$

which is the probability of cell $j'$ coming from subject $i$. The difference is that equation (14) depends on $X_{i'j'}$ while (15) doesn't. For later use, let $p_{i|x} = \mathbb{P}(S_{i'j'} = i \mid X_{i'} = x, \mathbf{X}, \boldsymbol{\alpha}_k)$ and $p_i = \mathbb{P}(S_{i'j'} = i \mid \mathbf{X}, \boldsymbol{\alpha}_k)$.

Let $M_0 = \sum_{i=1}^n \exp(\alpha_{ik}) p_{i|0}$, $M_1 = \sum_{i=1}^n \exp(\alpha_{ik}) p_{i|1}$, and $M = \sum_{i=1}^n \exp(\alpha_{ik}) p_i$. Substituting equation (14) and (15) to log of equation (13) gives

$$\begin{aligned} &\log \mathbb{E}\left[\exp\left(\alpha_{i'k}\right) \mid X_{i'j'}, \mathbf{X}, \boldsymbol{\alpha}_k\right] \\ &= \begin{cases} \left(\log M_1 - \log M_0\right) X_{i'j'} + \log M_0 & : \text{Scenario 1} \\ \log M & : \text{Scenario 2} \end{cases} \end{aligned} \tag{16}$$

Substituting equation (16) to equation (12) gives

$$\begin{aligned} &\mathbb{E}\left[Y_{i'j'k} \mid X_{i'j'}, \alpha_{i'k}, \mathbf{X}, \boldsymbol{\alpha}_k\right] \\ &= \exp\left(\beta_{0k} + X_{i'j'}\beta_{1k} + \log \mathbb{E}\left[\exp\left(\alpha_{i'k}\right) \mid X_{i'j'}, \mathbf{X}, \boldsymbol{\alpha}_k\right]\right) \\ &= \begin{cases} \exp\left(\log M_0 + \beta_{0k} + X_{i'j'}[\beta_{1k} + \log M_1 - \log M_0]\right) & : \text{Scenario 1} \\ \exp\left(\log M + \beta_{0k} + X_{i'j'}\beta_{1k}\right) : \text{Scenario 2} \end{cases} \end{aligned} \tag{17}$$

Thus, the expectation of the estimator $\widehat{\beta}_{1k}$ conditional on $\mathbf{X}$ and $\boldsymbol{\alpha}$ is

$$\mathbb{E}\left[\widehat{\beta}_{N,1k} \mid \mathbf{X}, \boldsymbol{\alpha}_k\right] = \begin{cases} \beta_{1k} + \log M_1 - \log M_0 & : \text{Scenario 1} \\ \beta_{1k} & : \text{Scenario 2} \end{cases} \tag{18}$$

because these are the coefficients of $X_{ij}$ in equation (17).

Subsequently, the variance is

$$\text{Var}\left(\mathbb{E}\left[\widehat{\beta}_{N,1k} \mid \mathbf{X}, \boldsymbol{\alpha}_k\right]\right) = \begin{cases} \text{Var}\left(\log M_1 - \log M_0\right) & : \text{Scenario 1} \\ 0 & : \text{Scenario 2} \end{cases} \tag{19}$$

The expansion of $\text{Var}\left(\log M_1 - \log M_0\right)$ gives

$$\begin{aligned} &\text{Var}\left(\log M_1 - \log M_0\right) \\ &= \text{Var}\left(\log M_1\right) + \text{Var}\left(\log M_0\right) \quad \text{(sampling of subjects is independent)} \\ &= \text{Var}\left(\log\left[\sum_{i:X_i=1} \exp(\alpha_{ik}) p_{i|1}\right]\right) + \text{Var}\left(\log\left[\sum_{i:X_i=0} \exp(\alpha_{ik}) p_{i|0}\right]\right) \end{aligned} \tag{20}$$

Since $p_{i|x} \sim 1/n$ for all subjects $i$ (it means that the number of cells from the subjects are in a similar order), the scale of $M_x$ $(x = 0, 1)$ is

$$\mathcal{O}(M_x) = \mathcal{O}\left(\frac{1}{n} \sum_{i:\text{all subjects}} \exp(\alpha_{ik})\right) = \mathcal{O}\left(\frac{1}{\sqrt{n}}\right) \tag{21}$$

by the central limit theorem applied to independent subjects. Therefore, the variance of the $\log M_x$ is $\mathcal{O}(1/n)$.

Now we arrive at our main conclusion.

$$\begin{aligned}
\operatorname{Var}\left(\widehat{\beta}_{N,1k}\right) &= \operatorname{Var}\left(\mathbb{E}\left[\widehat{\beta}_{N,1k} \mid \mathbf{X}, \boldsymbol{\alpha}_k\right]\right) + \mathbb{E}\left[\operatorname{Var}\left(\widehat{\beta}_{N,1k} \mid \mathbf{X}, \boldsymbol{\alpha}_k\right)\right] \\
&= \begin{cases} \mathcal{O}\left(\frac{1}{n}\right) + \mathcal{O}\left(\frac{1}{N}\right) & : \text{Scenario 1} \\ 0 + \mathcal{O}\left(\frac{1}{N}\right) & : \text{Scenario 2} \end{cases} \\
&= \begin{cases} \mathcal{O}\left(\frac{1}{n}\right) & : \text{Scenario 1} \\ \mathcal{O}\left(\frac{1}{N}\right) & : \text{Scenario 2} \end{cases}
\end{aligned} \tag{22}$$

⊠

**Theorem 1** clarifies the name of pseudoreplication bias. Virtually all tests, if not most, rely on the central limit theorem to test significance. A normal distribution approximation is applied at a certain step in the procedure to obtain $P$-values. Loosely speaking, the goodness of the approximation depends on the number of effective sample size, and this number is used to assess significance. The number also measures the amount of data available for statistical inference. In multi-subject scRNA-seq analysis, this number is ambiguous as there are at least two levels of units: the subjects and the cells. **Theorem 1** resolves the ambiguity by explicating that the number is the number of subjects ($n$) and the number of cells ($N$) in each design, respectively. Hence, cells are not the true replicates that drive the normal approximation of the test in the first scenario, so we call them pseudoreplicates. Nevertheless, cell-wise tests always take $N$ as the effective sample size, mistaking the true replicates as the cells. Therefore, it's called the pseudoreplication bias for using pseudoreplicates (cells) instead of true replicates (subjects) for the test.

### 4 Robust methods under dropouts

Let $Z_{ijk} = 0, 1$ is the dropout indicator variable that is 1 when dropout occurs and 0 when it doesn't. Then, a zero-inflated count $W_{ijk}$ is

$$W_{ijk} = Z_{ijk} \cdot 0 + (1 - Z_{ijk}) \cdot Y_{ijk} \tag{23}$$

where $Y_{ijk}$ follows the distribution that obeys the conditional mean equation (1).

Assuming that the dropout event occurs independent of $Y_{ijk}$ conditional on $\alpha_{ik}$ and $X_{ij}$,

$$\begin{aligned}\mathbb{E}\left[W_{ijk} \mid X_{ij}, \alpha_{ik}\right] &= \mathbb{E}\left[(1 - Z_{ijk})Y_{ijk} \mid X_{ij}, \alpha_{ik}\right] \\ &= \mathbb{E}\left[(1 - Z_{ijk}) \mid X_{ijk}, \alpha_{ik}\right] \cdot \mathbb{E}\left[Y_{ijk} \mid X_{ij}, \alpha_{ik}\right]\end{aligned} \tag{24}$$

which implies

$$\begin{aligned}\log \mathbb{E}\left[W_{ijk} \mid X_{ij}, \alpha_{ik}\right] &= \log \mathbb{E}\left[(1 - Z_{ijk}) \mid X_{ij}, \alpha_{ik}\right] + \log \mathbb{E}\left[Y_{ijk} \mid X_{ij}, \alpha_{ik}\right] \\ &= \log \mathbb{E}\left[(1 - Z_{ijk}) \mid X_{ij}, \alpha_{ik}\right] + \beta_{0k} + X_i\beta_{1k} + \alpha_{ik}\end{aligned} \tag{25}$$

Since $X_{ij}$ is discrete and the number of subjects is finite, $\log \mathbb{E}\left[(1 - Z_{ijk}) \mid X_{ij}, \alpha_{ik}\right]$ can be written as a linear function respect to $X_{ij}$ and $\alpha_j$ using dummy variables. Therefore, equation (25) obeys the conditional mean equation (1). Therefore, robust methods only requiring the conditional mean equation remain valid when dropout is present.
